# Whole-brain modeling explains the context-dependent effects of cholinergic neuromodulation

**DOI:** 10.1101/2022.03.21.485145

**Authors:** Carlos Coronel-Oliveros, Carsten Gießing, Vicente Medel, Rodrigo Cofré, Patricio Orio

**Affiliations:** Doctorado en Ciencias Mención Biofísica y Biología Computacional, Universidad de Valparaíso, Valparaíso, Chile; Latin American Health Brain Institute (BrainLat), Universidad Adolfo Ibáñez, Santiago, Chile; Centro Interdisciplinario de Neurociencia de Valparaíso (CINV), Universidad de Valparaíso, Valparaíso, Chile; Carl von Ossietzky University of Oldenburg, Department of Psychology, Oldenburg, Germany; University of Sydney, Brain and Mind Centre, Sydney, Australia; Universidad de Chile, Departamento de Neurociencia, Santiago, Chile; Institute of Neuroscience (NeuroPSI), Paris-Saclay University, Centre National de la Recherche Scientifique (CNRS), Gif-sur-Yvette, France; CIMFAV-Ingemat, Facultad de Ingeniería, Universidad de Valparaíso, Valparaíso, Chile; Instituto de Neurociencia, Facultad de Ciencias, Universidad de Valparaíso, Valparaíso, Chile

**Keywords:** Acetylcholine, fMRI, Neuromodulation, Nicotine, Segregation, Whole-brain models

## Abstract

Integration and segregation are two fundamental principles of brain organization. The brain manages the transitions and balance between different functional segregated or integrated states through neuromodulatory systems. Recently, computational and experimental studies suggest a pro-segregation effect of cholinergic neuromodulation. Here, we studied the effects of the cholinergic system on brain functional connectivity using both empirical fMRI data and computational modeling. First, we analyzed the effects of nicotine on functional connectivity and network topology in healthy subjects during resting-state conditions and during an attentional task. Then, we employed a whole-brain neural mass model interconnected using a human connectome to simulate the effects of nicotine and investigate causal mechanisms for these changes. The drug effect was modeled decreasing both the global coupling and local feedback inhibition parameters, consistent with the known cellular effects of acetylcholine. We found that nicotine incremented functional segregation in both empirical and simulated data, and the effects are context-dependent: observed during the task, but not in the resting state. In-task performance correlates with functional segregation, establishing a link between functional network topology and behavior. Furthermore, we found in the empirical data that the regional density of the nicotinic acetylcholine *α*_4_*β*_2_ correlates with the decrease in functional nodal strength by nicotine during the task. Our results confirm that cholinergic neuromodulation promotes functional segregation in a context-dependent fashion, and suggest that this segregation is suited for simple visual-attentional tasks.

## 1 Introduction

The flexible and adaptive behavior of vertebrate animals in different environmental contexts is partly possible due to the brain macroscale “anatomical architecture” also denoted Structural Connectivity (SC), which quantifies the existence of white matter tracts physically interconnecting pairs of brain regions. The brain SC sets the basis for the processing of information by domain-specific systems (segregation) and allows the binding of this specialized processing (integration) to guide adaptive behavior [1]. In fact, integration and segregation are proposed as prominent organizational principles in the brain [2, 3, 4], and their balance is a crucial element for the dynamical richness and flexibility of neural activity, necessary for the coherent global functioning of the brain [5, 6].

At the macro-scale, neural activity can be measured through an electroencephalogram (EEG) and functional magnetic resonance imaging (fMRI) recordings, among other techniques. The functional brain organization is typically characterized using pairwise statistical interdependencies between brain areas, building functional connectivity (FC) matrices [7, 8]. When applying graph theoretical approaches to the brain’s FC, striking fluctuations in the network topology appear between different conditions. In particular, a more segregated brain network is related to specific motor tasks, making brain regions more domain-specific [3]. In contrast, a reconfiguration towards the integration of the brain’s network is observed in complex tasks and arousal-related states [3, 9]. Interestingly, it has been shown that FC in resting-state (RS) conditions reveal a “default” balance between integration and segregation [3, 10, 11]. This balance changes toward segregation or integration as a function of behavioral tasks [3, 10], or by pathological conditions [12, 13, 14]. In contrast, the SC, which is the basis for shaping the dynamics of the brain, remains fixed during the short time scales relevant to cognitive tasks. Thus, how does the brain manage the transitions between different FC patterns, starting from a fixed connectome? A plausible candidate mechanism corresponds to neuromodulatory systems [4].

Neuromodulators change the neuron’s response profile without necessarily causing the cell to fire [15, 16], increasing or decreasing the probability of neurons to fire actions potentials. A recent hypothesis points out that the cholinergic and noradrenergic systems are key elements in promoting more segregated and integrated states, respectively [4]. At the behavioral level, acetylcholine is involved in several cognitive processes, such as attention, cue detection, and memory [17]. In the cortex, it has been suggested that acetylcholine could produce a shift from an internal-driven –high influence of excitatory feedback connections– to sensory-driven processing of information, enhancing the precision of sensory coding [18, 19]. Following this idea, the cholinergic system can reduce the influence of long-range feedback connections, through mechanisms that rely on both muscarinic and nicotinic receptors [18, 20, 21]. Also, the cholinergic system can increase the excitability of pyramidal neurons by reducing the somatic feedback inhibition mediated by interneurons [18, 22, 23]. Regarding nicotinic acetylcholine receptors, activation of *β*_2_-containing receptors by acetylcholine enhances the firing rates of dendritic-targeting gabaergic interneurons–for example, inhibitory Martinotti cells [24, 25]– increasing the inhibitory tone onto the apical section of deep layers pyramidal neurons, and interfering with the feedback connections that arrive to that section [18, 21, 26].

Computational and experimental works [9, 27, 28] suggest that the effects of noradrenergic and cholinergic neuromodulation are context-dependent, that is, different during tasks than in RS. In the case of noradrenaline, its role in promoting functional integration was previously addressed by [9, 10, 29]. At the whole-brain level, several mechanisms have been proposed to explain the effects of acetylcholine [15, 28, 30, 31]. However, there is still no clear relation between the context-dependent cholinergic modulation and large-scale network topology, as related to the cholinergic effect on promoting functional segregation [4]. A direct link between the empirical data, functional network topology, and computational modeling of the cholinergic system is missing. This constitutes the focus of the present work.

Here, we analyzed fMRI data previously published in [32], in both RS and task conditions (Go/No-Go attentional task), under the effects of nicotine. Specifically, we analyzed the FC using tools from graph theory to quantify integration, as global efficiency, and segregation, as transitivity [33]. Then, we built a whole-brain model from Jansen & Rit neural masses [34] interconnected using a human connectome. fMRI BOLD-like signals were generated from the firing rates of neural masses [35] and simulated FC matrices were computed. The model was fitted to reproduce the empirical FC matrices, minimizing the Euclidean distance between the simulated and empirical FC matrices in each condition (placebo and nicotine) and blocks (RS and task). We simulated the effects of nicotine decreasing the global coupling and feedback inhibition parameters in the model, as suggested previously [28]. The task was simulated by a heterogeneous change in the background input to brain regions. Our simulation results reproduce the context-dependent effects of nicotine, promoting functional segregation during the task, but not in RS.

At the behavioral level, we correlated the performance of the subjects in the Go/No-Go task with metrics computed from the empirical FC matrices. We found that segregation is a signature of in-task performance, showing a link between functional network topology and behavior. Finally, under the hypothesis that nicotine decreases global correlations, we verified if the regional density of cholinergic-related receptors obtained by positron emission tomography (PET) [36, 37, 38, 39, 40, 41] correlates with regional changes in FC. We observed that changes in nodal strength FC, during the task, correlate with the regional density of the *α*_4_*β*_2_ nicotinic acetylcholine receptor. However, the model with the receptor density map does not explain the effects of nicotine on FC better than a model that assumes uniform or random neuromodulation.

## 2 Methods

### 2.1 Ethics statement

Experimental data used in this work was previously collected in [32], where the ethical approval was obtained from the ethics committee of the German Psychological Association. Participants wrote informed consent for all procedures, which follow the principles of the Declaration of Helsinki.

### 2.2 Subjects and nicotine administration

The study involves a re-analysis of a previously published dataset [32]. Eighteen healthy right-handed cigarette smokers were studied in a double-blind, randomized, placebo-controlled, crossover design in which once nicotine was delivered in the form of a 4 mg nicotine polacrilex gum (Nicorette, McNeil AB) and once a placebo in the form of a taste- and size-matched gum.

The gum was chewed for 30 minutes; then the scanning started immediately. Subjects were asked to refrain from smoking 2 hours before the experiment.

### 2.3 Paradigm and task

The placebo and nicotine sessions were separated by at least 2 weeks and each session was divided into different time periods in which resting-state oscillations or task activations were measured using fMRI (Fig. S1). Three resting-state blocks comprising 256, 256, and 512 scans (6.4, 6.4, and 12.8 minutes, respectively) were measured alternately with two task blocks of 339 scans 8.5 minutes). During task blocks, participants performed a reversed version of the continuous performance test, measuring processes related to attention and vigilance [42]. During resting-state periods, participants were instructed to lie within the scanner, to keep their eyes open and to attend to a centrally presented visual stimulus.

### 2.4 fMRI data acquisition

Scanning was performed with a Siemens MAGNETOM Sonata MRI system (1.5 T). 1729 T2*-weighted echo-planar images with BOLD contrast were measured with a repetition time of 1.5 s and echo time of 50 ms. Detailed information on the investigated sample, nicotine administration, the paradigm, and the fMRI data acquisition can be found in [32].

### 2.5 Preprocessing and time series analysis

Preprocessing steps consisted in slice timing offsets correction, removal of head movements artifacts by alignment to the first volume, co-registration with the subjects’ anatomical images, spatial normalization to standard stereotaxic MNI space (Montreal Neurological Institute; http://www.mni.mcgill.ca/), and detrending. Next, nuisance covariates associated with the six linear head movements parameters, and anatomical masks for the cerebrospinal fluid (CSF) and white matter (WM) were regressed out from the data [43]. To remove the possible effects of small and transient head movements on FC, we quantified the framewise displacements (FD) and the root mean square of voxelwise percentage signal change (DVARS) of the preprocessed time series. In accordance with previously reported methods for “scrubbing” [44], we removed all frames of data with FD > 0.5 mm and/or DVARS > 0.5%.

Then, fMRI-BOLD time series were extracted using the Schaefer 100 cortical atlas parcellation [45] and band-pass filtered between 0.01 and 0.08 Hz [46] with a 3rd order Bessel filter. FC matrices were calculated as pairwise Pearson’s correlations, from which the subsequent analysis of functional network properties was performed.

Average volumetric PET images were collected for three different cholinergic receptors and transporters across 5 studies with open-dataset [40], which include nicotinic *α*_4_*β*_2_ [36], muscarinic *M*_1_ [41], and the vesicular acetylcholine transporter (VAChT) [37, 39, 40].

### 2.6 Whole-brain neural mass model

We simulated neuronal activity using the Jansen & Rit neural mass model [34, 49]. In this model, a brain area consists of three populations of neurons: pyramidal neurons, excitatory and inhibitory interneurons (Fig. 1A). The dynamical evolution of the three populations within the brain areas is modeled by two blocks each. The first block transforms the average pulse density in average postsynaptic membrane potential (which can be either excitatory or inhibitory) (Fig. 1B). This block, denominated post synaptic potential (PSP) block, is represented by an impulse response function. For the excitatory outputs:

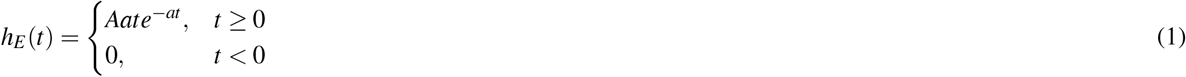

and for the inhibitory ones

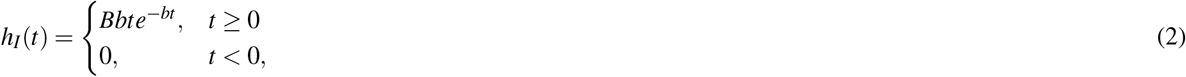

The constants *A* and *B* define the maximum amplitude of the PSPs for the excitatory (EPSPs) and inhibitory (IPSPs) cases, respectively, while *a* and *b* represent the inverse time constants for the excitatory and inhibitory postsynaptic action potentials, respectively. The second block transforms the postsynaptic membrane potential in average pulse density, and is given by a sigmoid function of the form

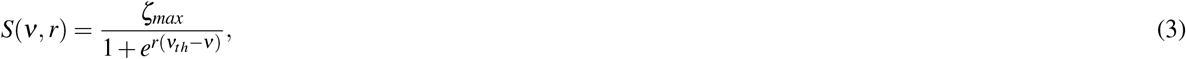

with *ζ_max_* as the maximum firing rate of the neuronal population, *r* the slope of the sigmoid function, *ν_th_* the half maximal response of the population, and *ν* their average PSP. Additionally, pyramidal neurons receive an external background input *p*(*t*), whose values were taken from a Gaussian distribution with mean *μ* = 5.4 impulses/s and standard deviation *σ* = 1. A parameter *β* heterogeneously modifies the background input in relation to the change in functional nodal strength 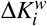 (see below, 2.9). In the model, each node *i* ∈ [1 … *n*] represents a single brain area. The global coupling is modified through a parameter *α*, and nodes are connected by a structural connectivity matrix *M* (Fig. 1C). This matrix was constructed using diffusion tensor imaging data (DTI) gathered from 100 unrelated subjects of the Human Connectome Project (HCP) 900 release [50], and parcellated in *n* = 100 cortical regions with the Schaefer 100 cortical atlas parcellation [45]; the SC is undirected and takes values between 0 and 1. The SC matrix was previously used in [51], and complete details of its construction can be found in the reference. Because long-range connections are mainly excitatory [52, 53], only links between the pyramidal neurons of a node *i* with pyramidal neurons of a node *j* are considered. The set of equations, for a node *i*, includes the within and between nodes activity

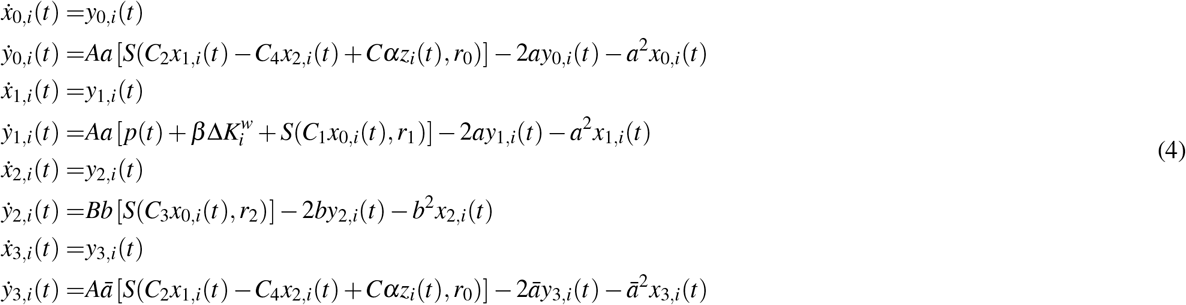

where *x*_0_, *x*_1_, *x*_2_ correspond to the local outputs of the PSP blocks of the pyramidal neurons, and excitatory and inhibitory loops, respectively, and *x*_3_ the long-range outputs of pyramidal neurons. The constants *C*_1_, *C*_2_, *C*_3_ and *C*_4_ scale the connectivity between the neural populations (see Fig. 1A). We used the original parameter values of Jansen & Rit [34, 49], except for *C*_4_: *ζ_max_* = 5 s^−1^, *ν_th_* = 6 mV, *r*_0_ = *r*_1_ = *r*_2_ = 0.56 mV^−1^, *a* = 100 s^−1^, *b* = 50 s^−1^, *A* = 3.25 mV, *B* = 22 mV, *C*_1_ = *C*, *C*_2_ = 0.8*C*, *C*_3_ = 0.25*C*, and *C* = 135. The parameter *C*_4_ was defined as a linear function of global coupling: *C*_4_ = (0.3 + 0.6*α*)*C*. The modification allowed the model to sustain oscillations in a wider range of *α* values, that is, it preserves the excitation/inhibition (E/I) balance. The term 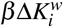 represents an additional heterogeneous input used to simulate the task block (see below). When simulating the RS block, *β* = 0.

**Figure 1.**
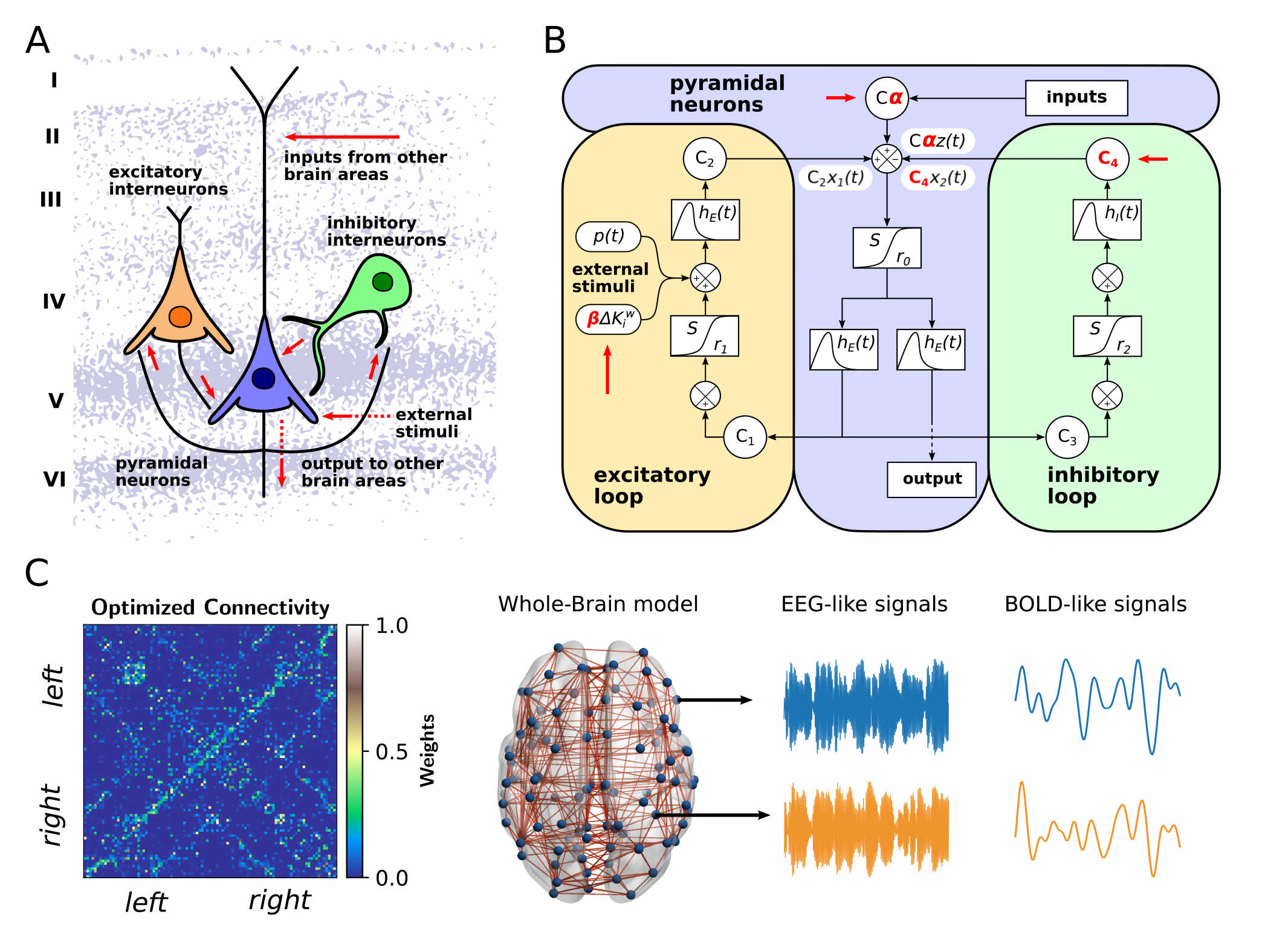
Model for whole-brain dynamics. **A)** Simplified scheme of the Jansen & Rit model, composed of a population of pyramidal neurons with excitatory and inhibitory local feedback mediated by interneurons. **B)** Overview of some important model parameters. Constants *C_i_* connect each population. The outputs are transformed from average pulse density to average postsynaptic membrane potential by an excitatory (inhibitory) impulse response function *h_E_*(*t*) (*h_I_*(*t*)). A sigmoid function *S* performs the inverse operation. Pyramidal neurons project to distant brain areas and receive both uncorrelated Gaussian-distributed inputs *p*(*t*) and inputs from other regions *z*(*t*), scaled by a global coupling parameter *α*. For simulating the task, a term 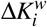 heterogeneously modifies the background input *p*(*t*) to pyramidal neurons. **C)** Each node represents a brain region, whose dynamics are ruled by the Jansen & Rit equations. The optimized connectivity matrix is the map of the connections (and their weights in the color bar) between brain regions (rows and columns of the matrix). This matrix was generated starting from the original human SC matrix. For each region, the model produces both EEG-like and BOLD-like signals. The brain figure in C) was obtained using the BrainNet Viewer [47]. The background figure in A) was adapted from [48] with permission.

The parameters *A*, *B*, *a* and *b* were selected to produce IPSPs longer in amplitude and latency in comparison with the EPSPs. The inverse of the characteristic time constant for the long-range EPSPs was defined as 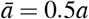. This choice was based on the fact that long-range excitatory inputs onto pyramidal neurons target their apical dendrites, and consequently this slows down the time course of the EPSPs at the soma [54].

The input from brain areas j ≠ *i* to the region *i* is given by

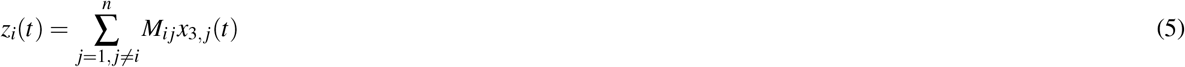

The average PSP of pyramidal neurons in region *i* characterizes the EEG-like signal in the source space; it is computed as [34, 49]

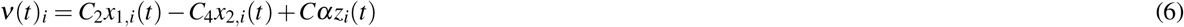

The firing rates of pyramidal neurons *ζ_i_*(*t*) = *S*(*ν*(*t*)_*i*_, *r*_0_) were used to simulate the fMRI BOLD-like signals.

### 2.7 Simulations

We ran simulations to generate the equivalent of 21 min real-time recordings, discarding the first 60 s. The system of stochastic differential equations (4) was solved with the Euler–Maruyama method, using an integration step of 1 ms. For model fitting and data augmentation (realizations of the fitted model), we used 100 and 300 random seeds, respectively, which controlled the stochasticity of the simulations. All the simulations were implemented in Python and the codes are freely available at https://github.com/vandal-uv/Nicotine-Whole-Brain.git.

### 2.8 Simulated fMRI-BOLD signals

We used the firing rates *ζ_i_*(*t*) to simulate BOLD-like signals using a generalized hemodynamic model presented in [35]. In this model, an increment in the firing rate *ζ_i_*(*t*) triggers a vasodilatory response *s_i_*, producing blood inflow *f_i_*, changes in the blood volume *v_i_* and deoxyhemoglobin content *q_i_*. The corresponding system of differential equations is

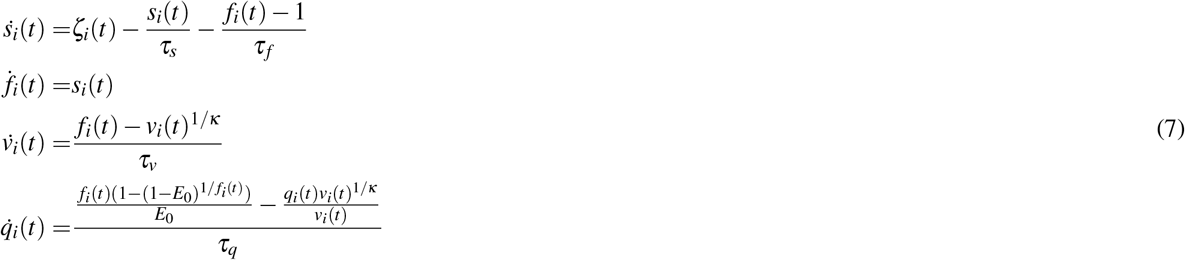

where *τ_s_* = 0.65, *τ_f_* = 0.41, *τ_v_* = 0.98, *τ_q_* = 0.98 represent the time constants for the signal decay, blood inflow, blood volume and deoxyhemoglobin content, respectively. The stiffness constant (resistance of the veins to blood flow) is given by *κ*, and the resting-state oxygen extraction rate by *E*_0_. We used *κ* = 0.32, *E*_0_ = 0.4. The BOLD-like signal of node *i*, denoted *B_i_*(*t*), is a non-linear function of *q_i_*(*t*) and *v_i_*(*t*)

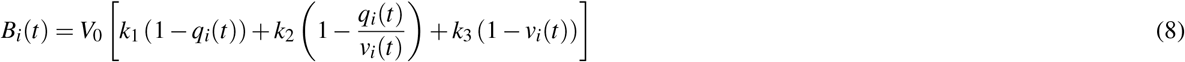

where *V*_0_ = 0.04 represents the fraction of venous blood (deoxygenated) in resting-state, and *k*_1_ = 2.77, *k*_2_ = 0.2, *k*_3_ = 0.5 are kinetic constants.

The system of differential equations (7) was solved using the Euler method with an integration step of 10 ms (firing rates were previously downsampled from 1000 Hz to 100 Hz). These BOLD-like signals were downsampled to the same sampling rate of empirical signals (TR = 1.5 s). Then, simulated signals were band-pass filtered between 0.01 and 0.08 Hz with a 3rd order Bessel filter, similar to the empirical data. These BOLD-like signals were employed to build the (FC) matrices –using pairwise Pearson’s correlations– from which the subsequent analysis of functional network properties was performed.

### 2.9 Nicotine and task modeling

To simulate the effects of nicotine, we decreased the value of the global coupling *α* and, consequently, the inhibitory-to-pyramidal neurons connectivity constant *C*_4_. This mimics the effect, in a simple way, of nicotinic and muscarinic acetylcholine receptors on long-range connectivity [18, 20, 21] and feedback inhibition [22, 23, 55]. For simulating the task block, we increase the factor *β*. This increases the mean background input that arrives at brain regions, in a manner that depends on the entries of 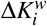; this term corresponds to the average empirical difference between the nodal FC strength in the RS and task blocks (placebo condition). The nodes that decrease their nodal strength the most, increment their background input the most [56]. The complete sequences of steps for model fitting are depicted in Fig. 3A.

### 2.10 Model fitting

We used the averaged empirical FC matrices across subjects as “targets” for model fitting. From the simulated BOLD-like signals, we built the simulated FC matrices with pairwise Pearson’s correlation. We vectorized the upper triangular of the empirical and simulated FC matrices, and then the vectors were compared using the Euclidean distance. A lower distance value corresponds to a better fit.

Before fitting the model to the empirical fMRI data, we performed an optimization procedure over the human SC matrix *M* [57, 58]. We swept the parameter *α* ∈ [0.5, 1.8], and chose the value of alpha that minimized the Euclidean distance of the empirical FC matrix in the placebo - RS block. This yielded *α* = 1.37 for an Euclidean distance and correlation, between matrices, around 15.4 and 0.23, respectively. From this point, we updated the values of the matrix *M* iteratively and discarded negative entries (negative values within the optimized SCs in each iteration were set to 0). The procedure was performed as follows:

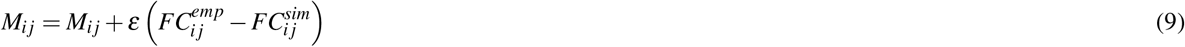

with 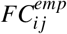 as the empirical FC matrix in the placebo-RS block, 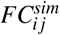 the simulated FC matrix at *α* = 1.37, and *ε* = 0.01 the rate of convergence. We used 64 random seeds in each iteration, and computed an averaged FC matrix across random seeds. The results of the optimization are presented in the Fig. S8, where the Euclidean distance decreases and correlation increases across iterations, indicating an improvement of the fitting. We employed this *optimized SC* matrix in the subsequent simulations, and it was not further changed. The codes for SC optimization can be found in https://github.com/vandal-uv/Nicotine-Whole-Brain.git.

The overall fitting is summarized in Fig. 3A. To fit the model to the placebo-RS and nicotine-RS blocks, we swept *α* ∈ [0.8, 1.5] in steps of 0.005 and searched for the minimum distance between FCs. Using the value of the placebo-RS block (*α* = 1.315), we swept the input slope *β* ∈ [0, 0.06] in steps of 0.001 for fitting the model to the placebo-task block. Then, employing *β* = 0.023 we swept *α* ∈ [0.8, 1.5] to fit the model to the nicotine-task block.

### 2.11 PET maps

We informed the model with acetylcholine-related receptors maps obtained by PET [36, 37, 38, 39, 40, 41]. Specifically, we used the nicotinic *α*_4_*β*_2_ receptor density map into Schaefer 100 parcellation. Also, we compared the original map with random and uniform versions of the former. Starting from the placebo condition (*α* = 1.315), we changed the *α* and *C*_4_ parameters linearly with the density vector of the *α*_4_*β*_2_ receptor maps, which was normalized to have an average mean equal to 1. Thus, we now have a different value of the global coupling, and also local feedback inhibition, for each node

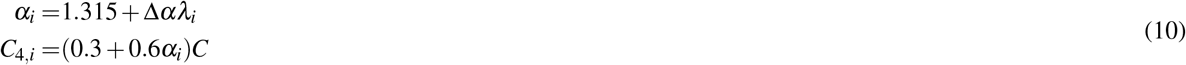

with Δ*α* as the increment in the regional *α_i_* parameter, and *λ_i_* the regional nodal value for the *α*_4_*β*_2_ receptor density vector. We obtained convex curves similar to the observed in the Fig. 3. A narrow range of Δ*α* values were employed: Δ*α* ∈ [0.04, 0.04] in steps of 0.002. We calculated the minimal Euclidean distance of the curves, and compared the values for each one of the maps employed. Before computing the minimal Euclidean distance, all the curves were smoothed using a 3 degree polynomial Savitzky-Golay filter and a window length of 17 points [59].

### 2.12 Functional integration and segregation

Starting from the FC matrices, we first applied a proportional threshold to remove spurious connectivity values [60]. Then, FC matrices were binarized and graph metrics were computed. Specifically, we calculated the binary global efficiency and transitivity, metrics related integration and segregation, respectively [33].

Global efficiency is based in paths, and was defined as [61]

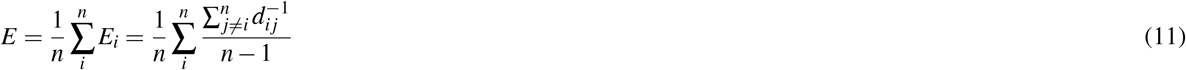

with *E_i_* as the nodal efficiency, *n* = 100 the total number of nodes, and *d_i j_* the shortest path that connects nodes *i* and *j*. The global efficiency is bounded between 0 and 1. Higher values are expected to be found in very integrated networks, where nodes can easily reach each other; values near 0 means the opposite. Transitivity is based in the count of the triangular motifs of the network, and it was computed as [62]

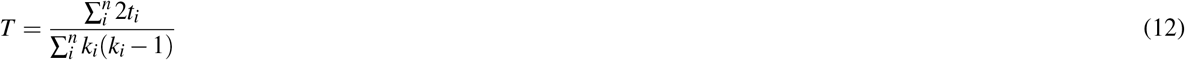

where *t_i_* corresponds to the number of triangles around the node *i*, and *k_i_* the node degree. The first one is defined as

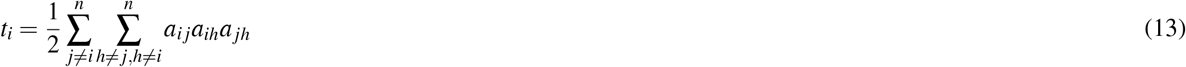

with *a_i j_* = 1 if nodes *i* and *j* are connected in the graph, and 0 otherwise. Transitivity captures the degree of local (short range) interactions between nodes and, in more common words, counts “in what extent my friends are also friends between them”. Higher values are expected from segregated networks, and 0 the opposite (e.g., completely random networks). Transitivity is a variant of the clustering coefficient, differing only on the normalization procedure: while transitivity is normalized collectively, clustering coefficient is normalized at the single node level [33].

To avoid the arbitrariness of choosing a single proportional threshold, we used a range of thresholds from 2 to 30% with linear increments of 1%. We report the area under the curve (AUC) of each graph metric in the threshold range explored [63]. The range of thresholds was set based on explorations, as a stable difference (nicotine - placebo) of *E* and *T* was found for network densities > 30%. Another issue regarding network analysis in FC is related to the influence of overall FC (the strength of the FC matrix) on functional network topology [60]. To divorce the effect of overall FC from topology, we regressed out the overall FC from each graph metric as suggested by [60]. Regression was done combining the subjects (or seeds) in all conditions (placebo and nicotine in both RS and task). The slope was used to regress out the overall FC, and finally we reported the corrected graph metrics. The graph analysis was performed using the Brain Connectivity Toolbox for Python (https://github.com/fiuneuro/brainconn) [33].

### 2.13 Hierarchical modular analysis of FC networks

To identify functional modules in both empirical and simulated data, we employed the hierarchical modular analysis (HMA) method following [64, 65]. Briefly, the method uses single value decomposition to find FC matrices’ eigenvectors and eigenvalues. Regions with the same signs within an eigenvector are assumed to have joint activity (cooperation), or the contrary when the signs are the opposite. The first functional level (first eigenvector) corresponded to the whole-brain network. The second level divides the brain in two modules, the third in four modules, and so on. For applying HMA, FC matrices must be symmetric and without negative entries. Also, the method does not require the use of thresholds. We divided each FC in 8 modules, and calculated the within and between modules connectivity. For subjects/seeds, the averaged within module connectivity (average across modules) is defined as

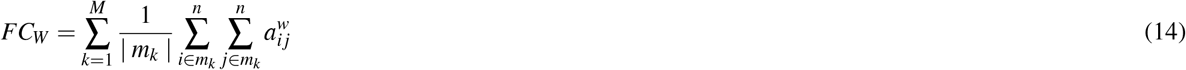

with *m_k_* being a module of the *M* = 8 total number of modules, and | *m_k_* | the number of links that belong to the module *m_k_*. Similarly, averaged between module connectivity is computed as

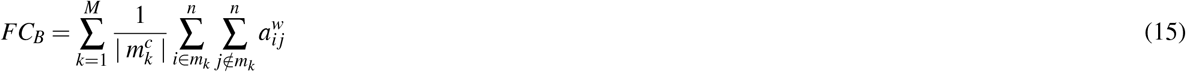

where 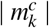 is the number of links that do not belong to the module *m*. Details of the method and calculations can be found in [64, 65]. We used our own implementation in Python (https://github.com/vandal-uv/Integration_Segregation_Methods) of the original codes of [65] (https://github.com/TobousRong/Hierarchical-module-analysis).

### 2.14 Network-based statistics

Beyond the global network analysis described in the last subsections, key connections that change under the effects of nicotine can be characterized using network-based statistics (NBS) [66]. The method identifies subsets of links that differ between two conditions and avoids the multiple comparisons problem when performing paired tests at the level of single links. We used the NBS implementation of the Brain Connectivity Toolbox for Python (https://github.com/fiuneuro/brainconn) [33, 66]. A one-tail paired t-test was used for ranking the connections, under the hypothesis that nicotine decreases functional connectivity (and then the strength of the links). A threshold of *t* = 3 was selected to obtain the connections that most change under the effects of nicotine. Next, the size of the largest component was calculated. Statistical significance of the found subnetworks was assessed with permutation tests, computing the size of the largest component for each permutation (up to 10000 permutations).

### 2.15 Statistical analysis

All pairwise comparisons, such as overall FC and network metrics, were performed with nonparametric permutation tests. These tests are suitable when using small sample sizes, and do not require any assumptions about normality [67]. The real difference between groups (computed as the mean difference) was compared with the distribution obtained from 10000 random surrogates; they were acquired by randomly reassigning the measures between groups (e.g., placebo and nicotine). We reported the *p* values, and results were considered as statistically significant for *p* values < 0.05. In addition, statistical relationships were measured using Pearson’s correlation.

To decrease the probability of making type I errors (false positives), we corrected all the *p*-values with the Benjamini-Hochberg method for multiple comparisons [68]. We applied the correction to paired comparisons (e.g., rest and task) and multiple correlations (e.g., reaction time vs FC metrics both in rest and task). In addition to using the *p*-values, which depend on statistical power, we computed the Cohen’s D distance (denoted using *D*) to report the results in terms of effect size, considering that the obtained *p*-values can be inflated by sample size computing additional realizations (model simulations) [69]. Depending on the absolute D value, effect sizes were classified as small, moderate, high, and huge for values greater than 0.2, 0.5, 0.8, and 1.2, respectively.

## 3 Results

### 3.1 Effects of nicotine on functional network topology are context-dependent

We first analyzed the empirical fMRI data [32], under the hypotheses that 1) nicotine promotes functional segregation, and 2) the effects are context-dependent. Recordings were taken from 18 healthy smokers in two different conditions: placebo, or under the effects of nicotine. BOLD-fMRI was measured in RS and during a Go/No-Go attentional task. The original data published in [32] comprises multiple RS and task periods (consecutive recordings). In this work, we principally focused our analysis and computational modeling on the fMRI and behavioral data of the first RS and task periods to avoid fatigue effects (see below).

There is a trend of nicotine to decrease the mean values within the FC matrices (referred to as overall FC, Fig. 2A); the effect was stronger in the task block than in RS. However, we did not find statistically significant differences in overall FC both in task (*p* = 0.1564, *D* = −0.36), and RS (*p* = 0.3238, *D* = −0.11). Then, the FC matrices were assessed for functional integration and segregation, as described in Methods. Segregation was measured by the transitivity *T*, while integration was quantified by the global efficiency *E*. As overall FC affects the thresholded FC network topology [60], we regressed out the overall FC from graph metrics with a simple linear regression (Fig. S2. Throughout this work, we report the corrected graph metrics.

**Figure 2.**
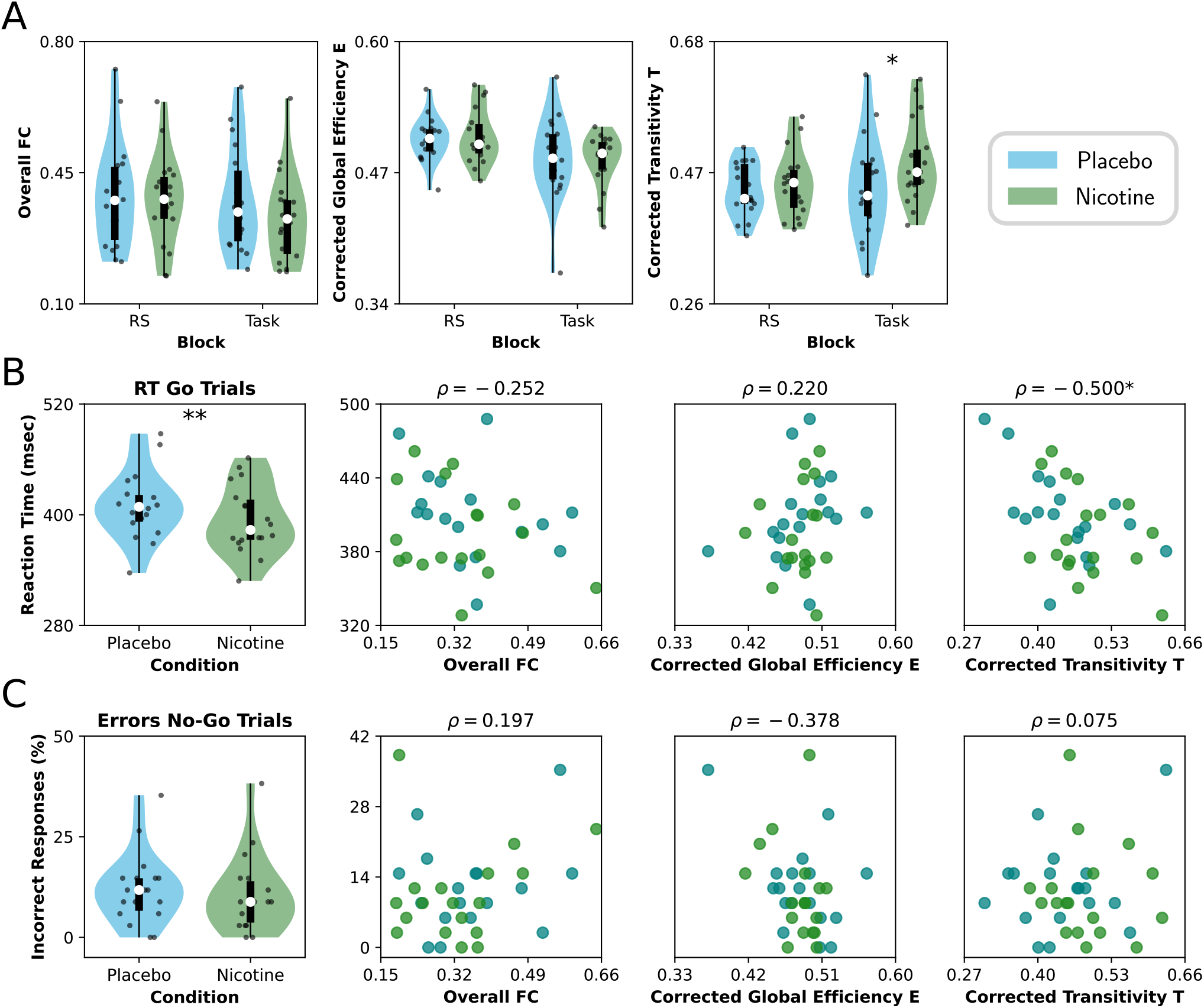
Empirical changes in functional connectivity produced by nicotine. **A)** Effect of nicotine on FC and functional network topology, both in RS and task. Violin plots represent the distributions of points (18 subjects per condition); the box plots represent the median (white circle), the first and third quartiles (box limits), and sample range (whiskers). **B)** Reaction time (RT) of the Go trials (left) and its correlation with FC metrics during the task. **C)** Percentage of incorrect responses of the No-Go trials (left) and correlation with FC metrics during task. Pearson’s *ρ* was used as a correlation measure. Points corresponded to participants (18 subjects per condition). ** : *p* < 0.01, * : *p* < 0.05.

**Figure 3.**
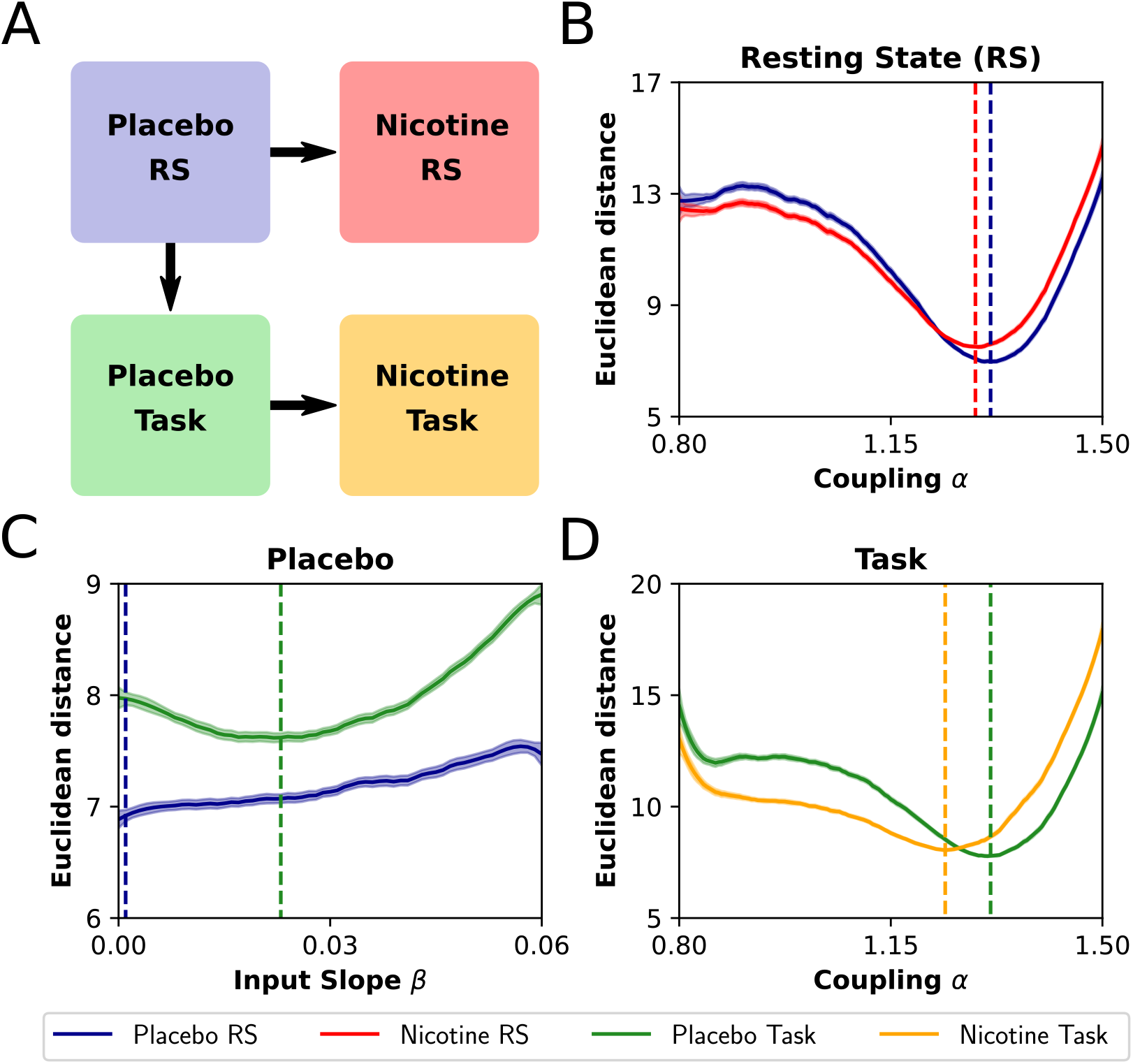
Fitting the whole-brain model. **A)** Starting from the model fitted to the placebo condition in RS (blue), the model was fit to the nicotine condition in RS sweeping the parameter *α* (red). The fit to the placebo condition during the task was performed by increasing *β* (green). Finally, Using the *β* value for the placebo-task, the model was fit for the nicotine-task by sweeping *α* (yellow). **B-D)** Euclidean distance between simulated and empirical FCs as a function of *α* and *β*. The minima of the Euclidean distances between empirical and simulated FC represent the values of the parameters *α* and *β* that better reproduce the empirical FCs. Segmented lines correspond to the optimal values found for placebo-RS (B and C) and placebo-task (D). A decrease in *α* was accompanied by a decrease in *C*_4_ (see text). Shaded areas represent the 95% confidence interval (using 100 random seeds).

Nicotine did not produce a change of *E* (integration) neither in RS (*p* = 0.4355, *D* = 0.07) nor during the task (*p* = 0.4355, *D* = −0.05) (Fig. 2A). On the contrary, nicotine increases *T* (segregation), being this effect statistically significant for the task block (*p* = 0.0438, *D* = 0.55), but not for RS (*p* = 0.1900, *D* = 0.2) (Fig. 2A). The result suggests that nicotine has a pro-segregation effect on the functional brain network topology, explained by the decrease in both overall FC and the increase in *T*. However, these effects were observed only in the task block, pointing out the context-dependent nature of nicotine modulation. Although we reported AUC as a summary measure of *E* and *T* across all thresholds, we show the results for each individual threshold in Fig. S3.

To characterize at a meso-scale level the effects of nicotine on functional network topology, we identified the key connections that change under nicotine in both RS and task using network based-statistics (NBS, see Methods) [66]. In RS, NBS showed that no links were significantly altered by nicotine (*p* = 0.7235). However, for the task block we found a subnetwork with links that decreased their functional strength under nicotine (*p* = 0.0421) (Fig. S4). To provide a quantitative measure, we computed the percentage of altered connections within and between the 7 RSNs (Fig. S4). We found that the connections between nodes of the visual, somatomotor, and dorsal attentional networks were the least affected by nicotine. Also, the default mode network reduces its connectivity with the rest of the brain networks, while the frontoparietal network preserves most of its connections with the visual, somatomotor, dorsal and ventral attentional networks. These findings suggest that nicotine reduces the influence of regions presumably not essential for the task, maintaining at the same time the connectivity profile of parieto-occipital and parieto-central regions.

To emphasize the context-dependent effect of nicotine, we repeated the network analysis on the three additional RS periods published in [32]. The first period is the one in Fig. 2A. The remaining periods were analyzed to explore the effect of fatigue on FC: time-on-task increases from RS period 1 to 4 (Fig. S5). Nicotine prevented the increase in overall FC triggered by time-on-task: across RS periods global correlations increase (as reported in the original article by [32]), and nicotine damped the increment in global correlations. Also, the increase of *E* and decrease of *T* caused by nicotine were more pronounced in RS periods 3 and 4, compared to periods 1 and 2. Thus, in conditions of prolonged time-on-task, nicotine dampens the changes in RS FC triggered by tiredness [32]. However, for the remaining analysis we will only include the first RS and task periods, because considering the effects of fatigue in our computational model falls beyond the scope of our work.

Next, to find a link between functional network topology and behavior (task performance), we analyzed the reaction time (RT) of the Go trials, the errors (percentage of incorrect responses) of the No-Go trials, and the metrics computed from the FC matrices in the task block. As previously reported [32], the main effect of nicotine was to decrease the RT of the subjects during task (*p* = 0.0037, *D* = −0.41) (Fig. 2B). Thus, subjects performed better under the effects of nicotine: a reduction in RT is a signature of enhanced attention [70]. When the RT is correlated with FC network metrics, we only found a significant negative correlation with the transitivity *T* (*p* = 0.0115), finding no correlation with the overall FC (*p* = 0.3960) and with *E* (*p* = 0.3960) (Fig. 2B). Regarding the error rate (Fig. 2C), there is a trend of nicotine to reduce the percentage of incorrect responses, but the result was not significant (*p* = 0.3208, *D* = −0.14). Also, we did not find a correlation between the percentage of incorrect responses with overall FC (*p* = 0.4980), *E* (*p* = 0.1384) and *T* (*p* = 0.6766). Thus, subjects that increase their segregation (transitivity) the most, also decrease their RT. From the results, it can be inferred that 1) a segregated network is suitable for the task (Go/No-Go), and 2) nicotine improves the reaction time of the subjects promoting a more segregated functional network during the task. To verify if the effects of the functional network topology on performance are specific to the task, we repeated the previous analysis using the metrics of the RS block (Fig. S6). We did not find a correlation between RT and errors with neither overall FC, *E*, nor *T* (*p* > 0.5 for all cases). Thus, RS metrics cannot predict the outcomes of the subjects in our Go/No-Go attentional task.

Finally, we checked if our results are independent of the parcellation considered. For that reason, we repeated the analyses presented in the (Fig. 2 using the automated anatomical labeling (AAL) parcellation [71], that divides the brain into 90 cortical and subcortical regions. With this parcellation (Fig. S7; *p* values and statistics in Tables S1 and S2), we found an increment of *T* only in the task block, and a negative correlation between the RT and *T*. This additional analysis supports the context-dependent effects of nicotine in promoting functional segregation, and its relevance to behavior (attention).

### 3.2 Modeling the effects of nicotine on functional connectivity

We simulated the whole-brain dynamics using the Jansen & Rit neural mass model [34] to find biophysical mechanisms for the effects of nicotine on functional network topology. As with the experimental data, brain areas were defined using the Schaefer 100 cortical parcellation, and connected with an SC matrix derived from DTI [51]. The model was also informed by the empirical fMRI BOLD signals [32] (see below).

The fitting steps are presented in Fig. 3A. First, using *β* = 0 (no task input) the model was fitted to RS, sweeping the global coupling *α* and comparing the simulated FC to an average of placebo and nicotine RS FCs. We found a minimum at *α* = 1.315, for placebo, and at *α* = 1.290, for nicotine (Fig. 3B). This confirms our previous hypothesis that a decrease in *α* and *C*_4_ can be used to mimic the effects of nicotine. Next, to simulate the effect of the placebo - task block, we swept the input slope *β* (Fig. 3C), finding a minimum at *β* = 0.023. A higher value of *β*, in general, represents an increase in the background input for almost all nodes, consistent with the idea that nodes that changed mostly their FC nodal strength, also increased their background input largely as an effect of the task [56]. Finally, using *β* = 0.023 we fitted the model to the nicotine - task block, sweeping *α* and finding a minimum at *α* = 1.240. Thus, a larger decrease in global coupling and feedback inhibition is required to simulate the effects of nicotine during the task.

Using the model with the fitted values of *α*, *C*_4_ and *β*, we obtained 300 simulated FCs (with different random seeds) for all four conditions (Fig. 4). Qualitatively, the averaged empirical (Fig. 4A) and simulated (Fig. 4B) FC matrices show that both nicotine and task decrease global correlations. Applying the network measures to the simulated FCs, our model reproduced the empirical results. First, nicotine decreases the overall FC (Fig. 4C) in both RS (*p* < 110^−5^, *D* = −0.59) and the task blocks (*p* < 110^−5^, *D* = −1.76), being the effect stronger in the latter. Similar to the empirical data, the following networks metrics were corrected for overall FC (Fig. S9). Second, we did not find an effect of nicotine on *E* both in task (*p* = 0.0723, *D* = −0.12) (Fig. 4C), and RS (*p* = 0.0566, *D* = −0.16). Lastly, nicotine increases *T* (Fig. 4C) and the effect was stronger and significant in the task block (*p* = 0.0032, *D* = 0.25) but not in RS (*p* = 0.3410, *D* = 0.03). Thus, both simulated and empirical data suggest that nicotine has a pro-segregation effect on network topology only in the task block.

**Figure 4.**
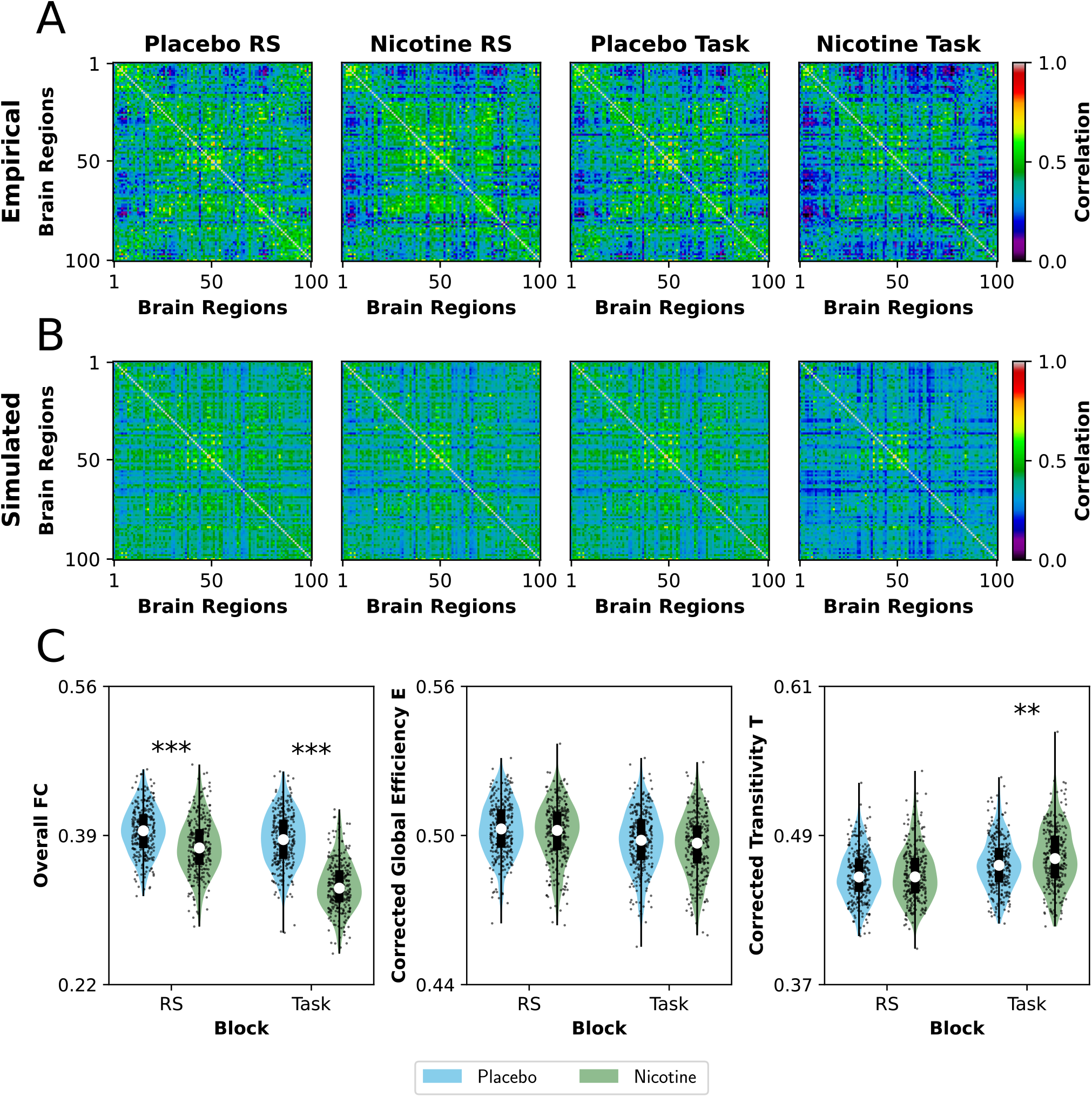
Simulated changes in functional connectivity produced by nicotine.. **A)** Empirical and **B)** simulated FC matrices, for all four conditions. For the empirical data, we show the average matrices across subjects (*n* = 18), and for the simulated data, we show the average using different random seeds (*n* = 300). **C)** Simulated effects of nicotine on FC and network topology, both in RS and task. Violin plots represent the distributions of points (300 random seeds per condition); the box plot is constituted by the median (white circle), the first and third quartiles (box limits), and sample range (whiskers). **: *p* < 0.01, *** : *p* < 0.001.

A possibility for explaining the context-dependent effect of nicotine on functional network topology is that, during the task, nicotine reduces more markedly the connectivity of distant regions in comparison to the more locally clustered ones. To test this possibility, we used Hierarchical Modular Analysis (HMA, see Methods) to group the nodes in 8 brain modules [64, 65], and finally reporting the between/within connectivity ratio. First, starting with the empirical data (Fig. 5B), we did not find any difference in the between/within connectivity ratio neither in RS (*p* = 0.3523, *D* = −0.09) nor task (*p* = 0.1761, *D* = −0.35)). However, in the simulated data, we observed that nicotine decreased the ratio in both conditions ([Fig. 5C]), the effect being stronger in the task block (*p* < 110^−5^, *D* = −1.36) than in RS (*p* < 110^−5^, *D* = −0.42). Thus, despite nicotine reducing global correlations, the effect in the task block is more pronounced on between-module connections than on within-module connections, explaining the shift to a more segregated functional network topology by nicotine during the task.

**Figure 5.**
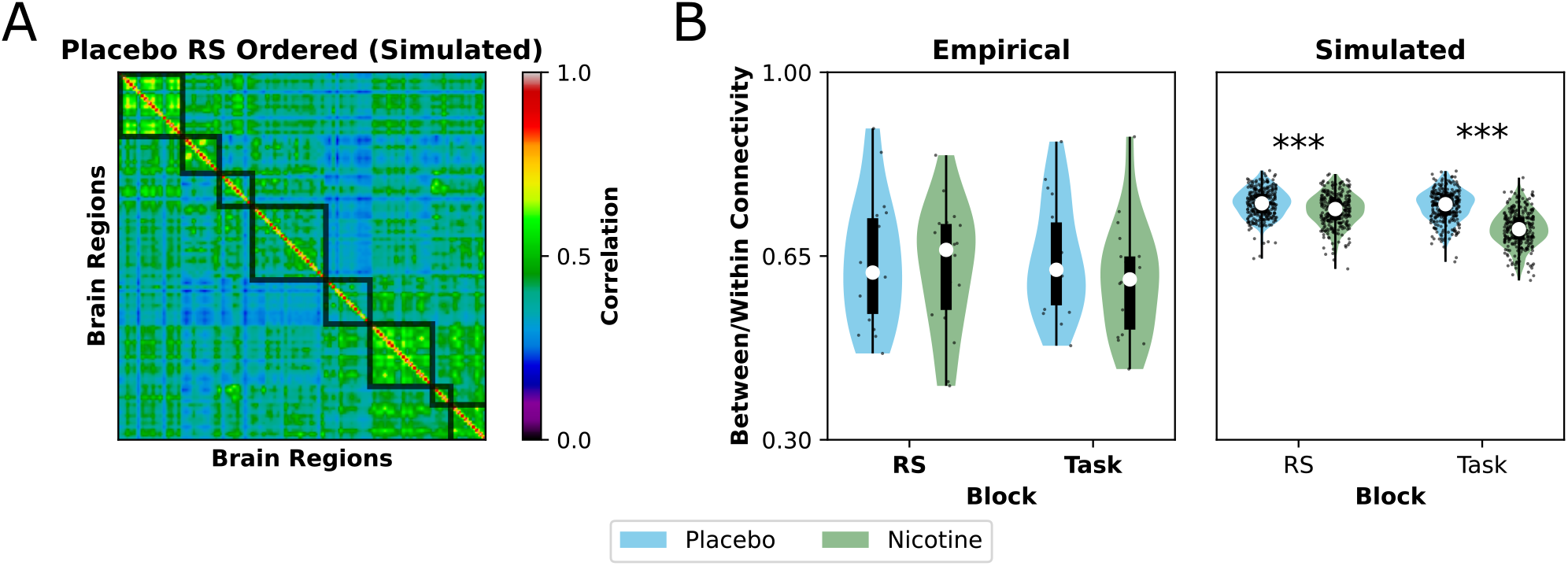
Modular connectivity under the effects of nicotine.. **A)** Example of the output of HMA on simulated data (averaged placebo RS simulated FCs). Nodes in FC matrix were ordered in accordance to their membership to 8 brain modules obtained using HMA. Between/within averaged connectivity ratio for **B)** empirical data (18 subjects) and **C)** simulated data (300 random seeds). Violin plots represent the distributions of points (18 subjects or 300 random seeds per condition); the box plot is constituted by the median (white circle), the first and third quartiles (box limits), and sample range (whiskers).*** : *p* < 0.001.

### 3.3 Nicotinic receptors partially mediate the changes in functional connectivity

Computational models suggest that the regional density of neurotransmitter receptors is a key element in reproducing the changes in brain dynamics by neuromodulation [72, 73]. To assess this possibility, we tested if the regional density of cholinergic-related receptors could explain the effects of nicotine on FC. We used regional receptor density maps, obtained with PET, of the *α*_4_*β*_2_ nicotinic receptor, the *M*_1_ muscarinic receptor, and the vesicular acetylcholine transporter (VAChT) [36, 37, 38, 39, 40, 41]. We calculated the difference in the empirical FC nodal strength between nicotine and placebo conditions (both in RS and task blocks) and correlated this vector with the density maps. In addition, we tested if the nodal strength of the optimized SC –a topological measure not related to receptors’ density– correlates with the effects of nicotine on FC.

Fig. 6 shows the correlation of SC nodal strength and receptor density with the changes in FC nodal strength. We did not find any correlation between the SC nodal strength and the change in FC neither in RS (*p* = 0.6723) nor task (*p* = 0.8636) (Fig. 6A). On the side of receptors, we only found in the task block a modest negative correlation between the density of the *α*_4_*β*_2_ receptor with the change in FC (*p* = 0.0471, Fig. 6A). Thus, these results suggest that 1) nodes expressing more *α*_4_*β*_2_ receptor tend to show a greater reduction of their functional nodal strength; 2) the *α*_4_*β*_2_ receptor plays a small role in producing the changes in FC during the task, but not in RS, and 3) SC cannot explain the specific (nodal) effects of nicotine on FC reduction. Considering the solid evidence that SC has a strong influence on the connectivity patterns observed in FC [74], it is striking to notice the absence of correlations between SC nodal strength with the changes in FC. For that reason, we computed the correlation of the FC nodal strength (in each condition and block independently) with the SC nodal strength (Fig S8). In the four conditions, there was a statistically significant positive correlation between the variables, especially in the placebo - RS block (Fig. S10). Although SC can influence FC, the changes observed under the effects of nicotine are more likely to be linked to the *α*_4_*β*_2_ receptor density.

**Figure 6.**
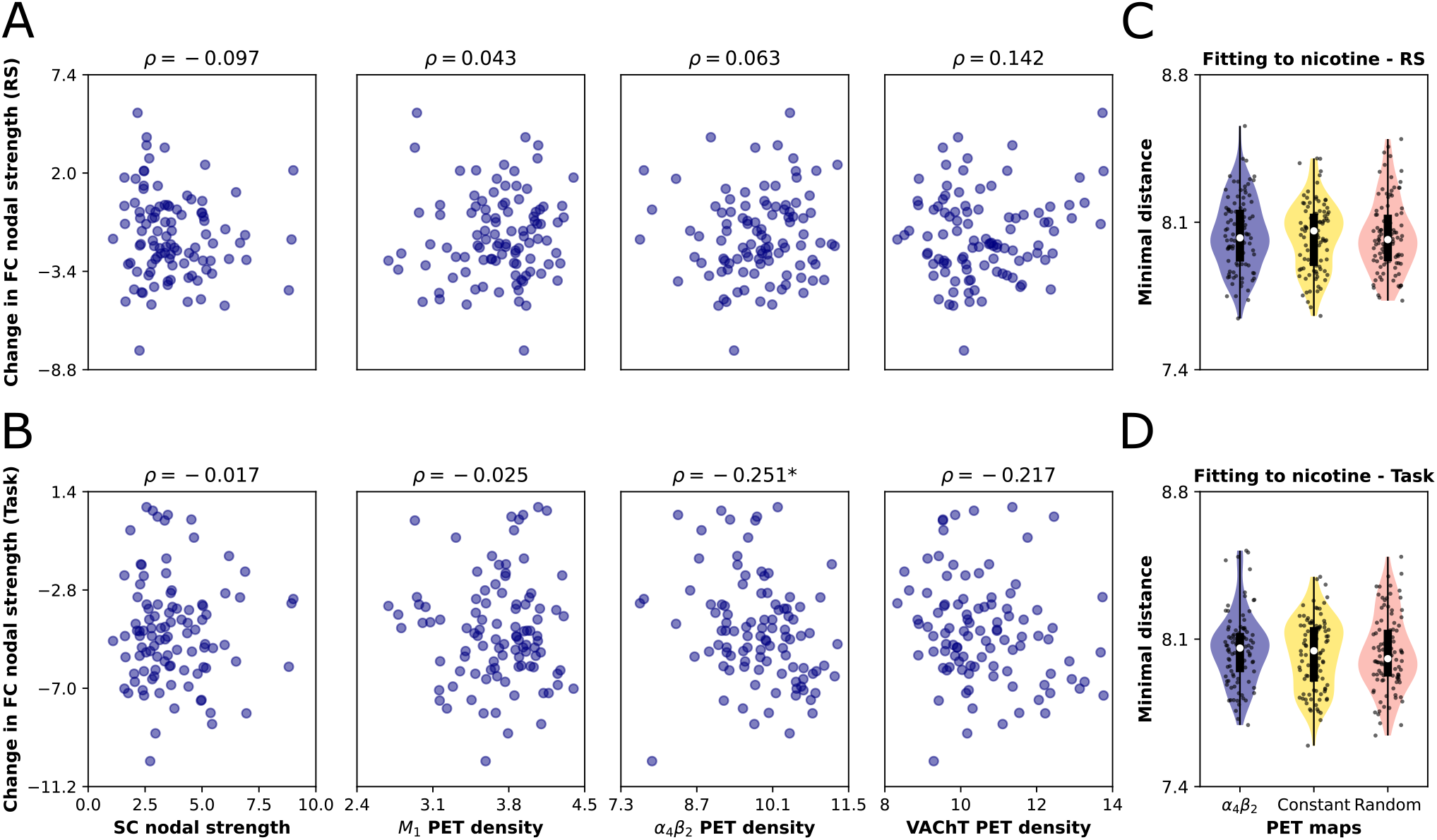
Relationship between cholinergic receptors and the nicotine-mediated effects on FC. **A-B)** Correlation between the change in empirical FC nodal strength (RS and task) with SC nodal strength and PET density of cholinergic-related receptors and transporters. **C-D)** Fitting to nicotine condition, both RS and task blocks, using the *α*_4_*β*_2_ receptor density map. We compared the fitting of the original map with two different surrogates (random and uniform maps derived from the original *α*_4_*β*_2_ receptor density vector). The minimal Euclidean distance is shown. Violin plots represent the distributions of points (300 random seeds per condition); the box plot is constituted by the median (white circle), the first and third quartiles (box limits), and sample range (whiskers). Points correspond to the Schaefer 100 brain regions. * : *p* < 0.05.

If the *α*_4_*β*_2_ receptor density correlates with the changes in FC, a model including the distribution of this receptor across the brain should improve the fitting to the empirical data. We analyzed this possibility by repeating the fitting performed in Fig. 3 for the nicotine condition, both in RS and task blocks. However, this time we changed the parameters *α* and *C*_4_ proportionally to the *α*_4_*β*_2_ receptor density, starting from the placebo condition (*α* = 1.315). Also, we used two different surrogates for comparison purposes. They consisted of randomization of the *α*_4_*β*_2_ receptor density vector, and the uniform vector used in Fig. 3. We did not find any improvement in model fitting when using the *α*_4_*β*_2_ receptor density map both in the task block (*p* = 0.4291, *D* = 0.01 and *p* = 0.2367, *D* = 0.06 for uniform and random maps, respectively) and in RS (*p* = 0.4291, *D* = −0.02 and *p* = 0.2367, *D* = 0.07, for uniform and random maps, respectively) (Fig. 6C-D). Consequently, if the *α*_4_*β*_2_ receptor plays a key –but partial – role in explaining the effects of nicotine on whole-brain FC, our model is not capable to capture the regional effects of the receptor.

## 4 Discussion

We have explored the effects of nicotine on the brain’s FC network topology by analyzing fMRI data and computational modeling. Our main results support the hypothesis that cholinergic neuromodulation promotes functional segregation [4] in a context-dependent fashion: the effect is more pronounced when performing a task than during resting-state. In addition, functional segregation is correlated with in-task performance (reaction time), suggesting a link between FC network topology and behavior. Our computational modeling supports the idea that the global effects on the FC network are the result of well-known local effects of acetylcholine: an increase in inhibition onto pyramidal neurons apical dendrites, alongside the decrease in somatic feedback inhibition. Finally, the empirical results posit the *α*_4_*β*_2_ receptor as an influential factor determining the effects of nicotine. However, informing our computational model with the *α*_4_*β*_2_ receptor density map did not improve the explaining capability of the model, a limitation we addressed below.

The stronger effect of nicotine on global correlations during the task may have several explanations. If we expect that the task by itself promotes the release of endogenous acetylcholine [75], then nicotine could amplify the effect of cholinergic modulation by directly binding to nicotinic receptors or by promoting the release of more acetylcholine [76]. In our computational model, the heterogeneous increase in background input –to mimic the effects of the task– acts as a mechanism of network control [77], driving the network to a more functional segregated state without any change in the brain SC. From this point, nicotine produces a higher contrast with the placebo condition, reflected in the reduction of integration and increase in segregation observed with nicotine. Further, our model suggests that the context-dependent effect of nicotine might be a consequence of a selective decrease of correlations, with between-module connectivity more affected than within-module connectivity (Fig. 5). A different fitting procedure could consist on start from the *α* value found for nicotine in RS condition, and then sweep *β* to find the best fit to the Nicotine-task condition. *β* should reduce, at least, the global correlations, similar to decreasing *α*. However, we would have faced a mechanistic election problem: is the shift to functional segregation by nicotine during task a consequence of the interaction between nicotine and task, triggering the release of more acetylcholine (the equivalent of increasing *α*)? Or, might nicotine have an influence in enhancing the task-related external stimuli (the equivalent of increasing *β*)? It is very possible that both *α* and *β* are modulated by the task and nicotine [18], but proving this hypothesis is beyond the scope of this work.

In our analysis, the degree of segregation correlates with in-task performance (Fig. 2B). In contrast, it has been well established that integration is associated with better performance in a wide range of behavioral tasks [78]. However, segregation might be a convenient feature in relatively simpler, more monotonous, or “automatic” motor tasks [78]. This was proposed by [3], arguing that simpler tasks require the coordination of a few brain subnetworks, and in this case, a segregated network topology should be desirable for optimal in-task performance. For example, enhanced segregation was reported in motor learning [79], visual-attentional [65] and sustained attention tasks [80]. In the specific case of [80], segregation increases in task-related networks (e.g. salience, cingulo-opercular, dorsal attention, and visual networks), while integration jumps up in brain regions not involved in the task (auditory network). The increase in segregation may prevent cross-interference between brain networks, allowing accurate processing of task-relevant information within the brain. On the other hand, the local increment of integration could constitute a mechanism of top-down suppression, through network integration, of regions not relevant for the task [80]. The role of the cholinergic system in selectively decoupling brain regions, to process in a more efficient way external stimuli, corresponds to the cholinergic shift to functional segregation during task [4].

Considering the effect of acetylcholine in enhancing attention in visual-spatial tasks [15, 18, 81], it is plausible to think that acetylcholine promotes segregation by increasing the within- and between-network connectivity in regions involved in visual processing and goal-directed attention. Our exploratory analysis using NBS is in line with the task-triggered increase in segregation reported by [80], where the increment in segregation was observed in regions involved in visual processing and attention. Also we observed a “disengagement” (links with reduced connectivity) of the frontoparietal network with default mode network (Fig. S4). The dorsal attentional network is known to play a role in the top-down voluntary guided allocation of attention [82], while the default mode network is more active when the subject is not focused in the external world [83]. The reduced connectivity between frontoparietal network with dorsal attentional network could constitute a shift from an internal- to external-driven processing of the information within the brain [84]. Furthermore, our results resonate with the work of [85]. Experienced players in real-time strategy video games, presented changes in brain SC that comprise an increase of within and between connectivity in occipital and parietal brain regions, both required for processing visual information. Topologically, they found an increase in local efficiency (segregation) within the occipito-parietal network. This case is interesting because the morph in network topology, which is non-transient (long-standing), might comprise plasticity changes in the SC that enhance the bandwidth of communication between and within occipital and parietal regions [85]. The cholinergic system could play a similar role in mediating transient changes in network topology, changes that facilitate the processing of visual information from noisy signals.

We found a negative correlation between the nicotinic *α*_4_*β*_2_ receptor density, obtained by PET, and the change in FC nodal strength. This result should not be a surprise, considering that FC changes were analyzed under the effects of nicotine. Despite this, the result emphasizes that the changes observed in FC can be ascribed to the cholinergic system and, more specifically, to nicotinic receptors. The nicotinic *α*_4_*β*_2_ receptor has a prominent role in attention and cognition [17]. For example, a dysfunction in *α*_4_*β*_2_ receptors is associated with cognitive impairments in mild Alzheimer’s dementia [86]. Also, stimulation of *α*_4_*β*_2_ receptors enhances performance in visual attentional tasks [87, 88], while their selective inhibition produces the opposite results [87]. This highlights the role of *α*_4_*β*_2_ receptor in mediating the nicotine-induced changes in FC. However, incorporating the *α*_4_*β*_2_ PET density map in the whole-brain model did not improve the fit to the nicotine condition both in RS and during the task. This constitutes a limitation of our model, and there are some possible explanations for those results. First, the changes may be small –as evidenced by the significant but relatively low correlation of FC nodal strength difference with the *α*_4_*β*_2_ receptor density map. Consequently, the model may not be able to capture the subtle functional re-configurations of the brain networks triggered by nicotine. Second, some theoretical works argue that the nodal properties, derived from the SC, can better explain the effects of neuromodulation on brain dynamics [29, 89, 90]. Considering this, it is plausible to think that the interaction between SC, neuromodulatory systems (and their receptors) and FC is not straightforward. Finally and perhaps more importantly, the PET maps used in our work studies are not subject-specific, a point that becomes more relevant when considering that the subjects were healthy smokers. There is evidence that nicotine can up-regulate nicotinic receptors [91, 92], and consequently, the regional distribution of the *α*_4_*β*_2_ receptor in the brain can diverge between healthy non-smokers and healthy smokers.

The mechanisms employed here are compatible with previous computational models, which attempted to simulate the effects of acetylcholine on global brain dynamics [28, 30, 93]. In these works, a decrease in global coupling and spike-frequency adaptation has been proposed to simulate the influence of acetylcholine in the transition from sleep to wakefulness [30, 93]. A similar mechanism was proposed by [28], who simulated the effects of acetylcholine by increasing the neural gain (the slope of the input-output function of neural masses) and decreasing global coupling. The main difference in our model was the introduction of a reduced feedback inhibition as an effect of nicotine, in contrast to the increment in neural gain used in [28]. However, in all the aforementioned cases (including our work), the increase in segregation is compatible with the reduction in global coupling by the cholinergic system. Indeed, the mechanisms are related to the shift from internal to external processing of information within the brain by acetylcholine [19], as a consequence of reducing the influence of feedback connections about external stimulation.

Another interesting finding is the lack of correlation between overall FC and task performance as measured by the RT, and in parallel, the negative correlation we found between segregation and RT. Although one of the main effects of nicotine in light of our results was to decrease the global correlations (overall FC), the change in network topology better-predicted subjects’ performance in the Go/No-Go task. Moreover, the topology of the FC network during the task was better at predicting performance during the task, in contrast to the topology of the FC network during RS. Thus, our findings suggest that 1) network topology matters for predicting performance during the task, and 2) as pointed out by [28], a characterization of RS FC is not sufficient to explain the effects of neuromodulatory systems on behavior. Following this idea, [9] reported a context-dependent effect of noradrenergic neuromodulation. The authors found an increase in RS segregation triggered by atomoxetine (a noradrenaline reuptake inhibitor). However, the opposite results were observed during an N-back task where atomoxetine increases functional integration. They suggest that the cognitive load of the N-back task, compared with the lack of cognitive constraints at RS, enhances phasic firing patterns of the locus coeruleus, which in turn could be potentiated by atomoxetine and ultimately promote functional brain integration. Similar results were found by [27], who analyzed the effects of clonidine (an *α*_2_-adrenergic autoreceptor agonist) on fMRI, reporting an increment of functional connectivity during the task and a decrease during RS.

Several ideas can be explored based on this work. For example, the mechanisms used here could be adapted to simulate the effects of the cholinergic system during the transition from sleep to wakefulness, similar to [30] and [93]. In addition, whole-brain models could be used to test specific targets for pharmacological treatments and to evaluate structure–function relationships [72, 94]. By combining fMRI, structural, and neurotransmitter PET data, personalized whole-brain models can be used to find specific targets for brain disorders [95]. This could lead to new treatments for disorders in which the cholinergic system is impaired, such as Alzheimer’s disease and age-related cognitive decline [96]. With a more theoretical motivation, we consider here only pairwise interactions and neglect higher-order effects that might contain relevant information related to high-dimensional functional brain interactions. Information theory [97], algebraic topological approaches [98, 99] and metaconnectivity [100] may provide useful information about high-order interdependencies in the brain. Indeed, a whole-brain model was proposed as a non-linear model of neurodegeneration to simulate the ageing of connectomes [101]. This model successfully reproduced the changes of high-order statistics observed in neuroimaging data [102], specifically, a significant increase in redundancy-dominated interdependencies between BOLD activity in older subjects. In this scenario, it would be interesting to reverse the ageing-triggered increment in redundancy by simulating the effect of cholinergic neuromodulation in a whole-brain model. On the other hand, the analysis performed here considers only the static (time-averaged) FC. How FC and functional network topology change over time should be addressed in future [10, 29, 103].

We must point out some limitations of our work. First, nicotine is not a full pro-cholinergic drug, as are donepezil, galantamine, and rivastigmine, which are cholinesterase inhibitors. However, nicotine may promote the release of acetylcholine [76], which is plausible given that nicotinic receptors are widely expressed in several types of neurons, including cholinergic neurons [104]. Moreover, the decrease in global correlations by nicotine reported here was also observed under the effects of donepezil [28]. In view of these points, we believe that our generalization, that is, linking the influence of nicotine to the pro-segregation effects of the cholinergic system [4], constitutes a reasonable statement. As a second limitation, the Jansen & Rit model (as other biophysically inspired models used at the whole-brain scale) is too complex to answer in a rigorous causal and mechanistic way (beyond correlations) why task activity promotes segregation. More simpler models and direct fMRI BOLD generative models ([57] as an example) can be used to explored the effects of perturbations (stimulation) on functional network topology. Finally, the empirical SC and the PET density maps are not subject-specific and, moreover, were obtained from different populations. We partially solved this problem using SC optimization, similar to [57, 58]. However, the unavailability of subject-specific empirical priors limited our work to simulate the effects of nicotine using an averaged observable of brain dynamics, i.e., the averaged FCs across subjects. We are aware of the potential of personalized whole-brain models for characterizing brain dynamics [95]. They can be used in clinical and translational neuroscience to propose model-inspired therapies, at the single subject level, with the aim to restore the normal brain function, and also to establish causal links between the model mechanisms (e.g., biophysical parameters fitted for individual patients) and the functional disturbances or cognitive decline ascribed to brain disorders [105, 106]. In spite of this, we believe that using individual whole-brain models might be not necessary for finding global principles of how context-dependent effects of cholinergic neuromodulation influence the functional network topology of the brain, as explored in our work. Also, we are aware of the relatively small experimental sample size used in this work. The lack of significant differences in empirical data –a possible consequence of the small sample size– limited the reach of HMA results, and consequently these results should be interpreted with caution. There is also strong evidence that fMRI measurements have poor precision/reliability and low power. In fact, one study claims that thousands of subjects are needed to show a reproducible association between individual differences in behavior and brain states [107]. In contrast to the majority of fMRI studies, our analysis is strongly theory-driven, which is helpful in improving the knowledge in the field and also making better use of small data sets [108].

### 4.1 Conclusions

Overall, our results suggest mechanisms to explain the pro-segregation effects of the cholinergic system, mechanisms that are compatible with previous computational models and the biophysics of nicotine action. We observed that the pro-segregation effects of nicotine, at the whole-brain level, depend on the behavioral context that subjects are experiencing. Furthermore, the degree of in-task segregation, captured as network transitivity, correlates with behavioral performance in a visual-attentional task. We suggest that, in this specific scenario, network segregation might be suitable for optimal in-task performance (segregation reduces reaction time). Finally, we observed that the regional expression of the nicotinic *α*_4_*β*_2_ receptor correlates with the changes in the functional nodal strength presented in empirical data, but only during the task block and not in RS. Including PET density maps into the model has no effects on the goodness of fit, pointing out that other factors might mediate the effects of nicotine on whole-brain functional connectivity.

To our knowledge, this is the first work that combines empirical fMRI data, in-task performance, and computational modeling to study the effects of the cholinergic system on functional network topology. Establishing links between neuromodulation and its functional correlates has a great scientific potential to deepen our understanding of the brain and its context-dependent response, and more generally to shed some light on the complex relationships between the structure and function of the human brain.

## Acknowledgements

We thank Gustavo Deco and Andrea Luppi for kindly providing the structural connectivity matrices used in the model. This work was supported by Fondecyt Grants 1211750 and the Advanced Center for Electrical and Electronic Engineering (FB0008 ANID, Chile) (PO), and the Human Brain Project, H2020-945539 (RC). The Centro Interdisciplinario de Neurociencia de Valparaíso (CINV) is supported by ICM-ANID, Grant ACE210014. CC-O is funded by Beca Doctorado Nacional ANID 2018-21180995, and the Latin American Brain Health Institute (BrainLat). The analyzed data derive from a study of [32] funded by Deutsche Forschungsgemeinschaft Grant GI 682/2-1 (CG).

## Author contributions statement

**Carlos Coronel-Oliveros**: Conceptualization, Investigation, Methodology, Formal analysis, Writing - original draft, Visualization, Validation. **Carsten Gießing**: Investigation, Data curation, Writing - review & editing, Supervision, Funding acquisition. **Vicente Medel**: Investigation, Methodology, Writing - review & editing. **Rodrigo Cofré**: Conceptualization, Investigation, Writing - review & editing, Methodology, Supervision. **Patricio Orio**: Conceptualization, Investigation, Writing - review & editing, Methodology, Supervision, Project administration, Resources.

## Data availability statement

All the codes used to perform the simulations are freely available in the next github repository https://github.com/vandal-uv/Nicotine-Whole-Brain. The graph analysis was performed using the Brain Connectivity Toolbox for Python (https://github.com/fiuneuro/brainconn) [33]. Codes for HMA can be found in https://github.com/vandal-uv/Integration_Segregation_Methods. Data are available on request from Dr. Carsten Gießing, University of Oldenburg. A formal data sharing agreement is required.

