## Supplementary Figures for "Whole-brain modeling explains the context-dependent effects of cholinergic neuromodulation"

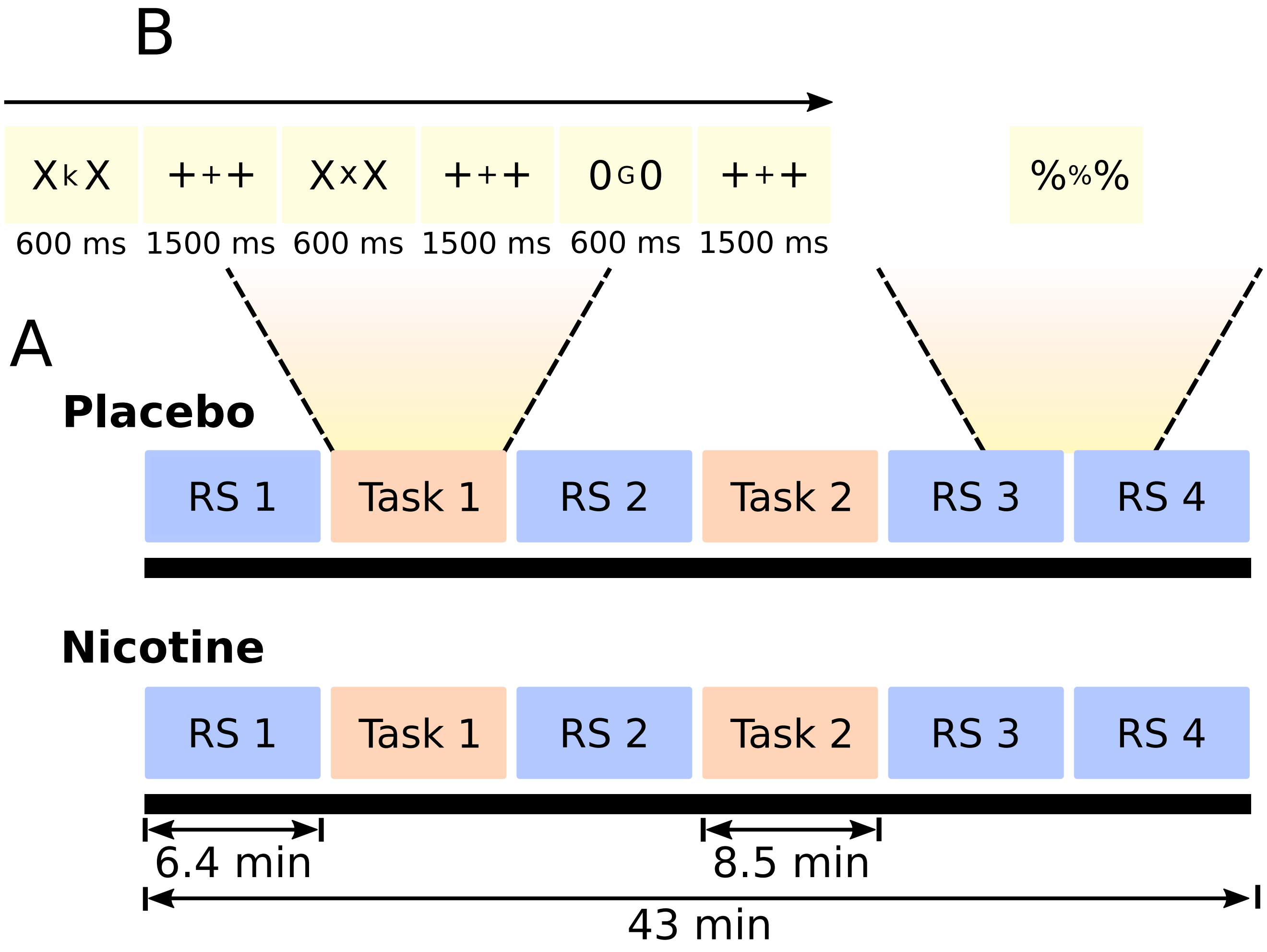


**Fig. S 1. Paradigm and task. A)** fMRI BOLD recordings were taken in several consecutive RS and task blocks. Two conditions were considered, placebo and nicotine, separated by at least two weeks. RS and task blocks had a duration of 6.4 and 8.5 minutes, respectively, for a total scan time of 43 min. **B)** Reversed version of the continuous performance test (Go/No-Go task). During RS periods, subjects kept their eyes open and looked at a centrally presented stimulus (%%%). Figure adapted from Gießing et al., (2013).

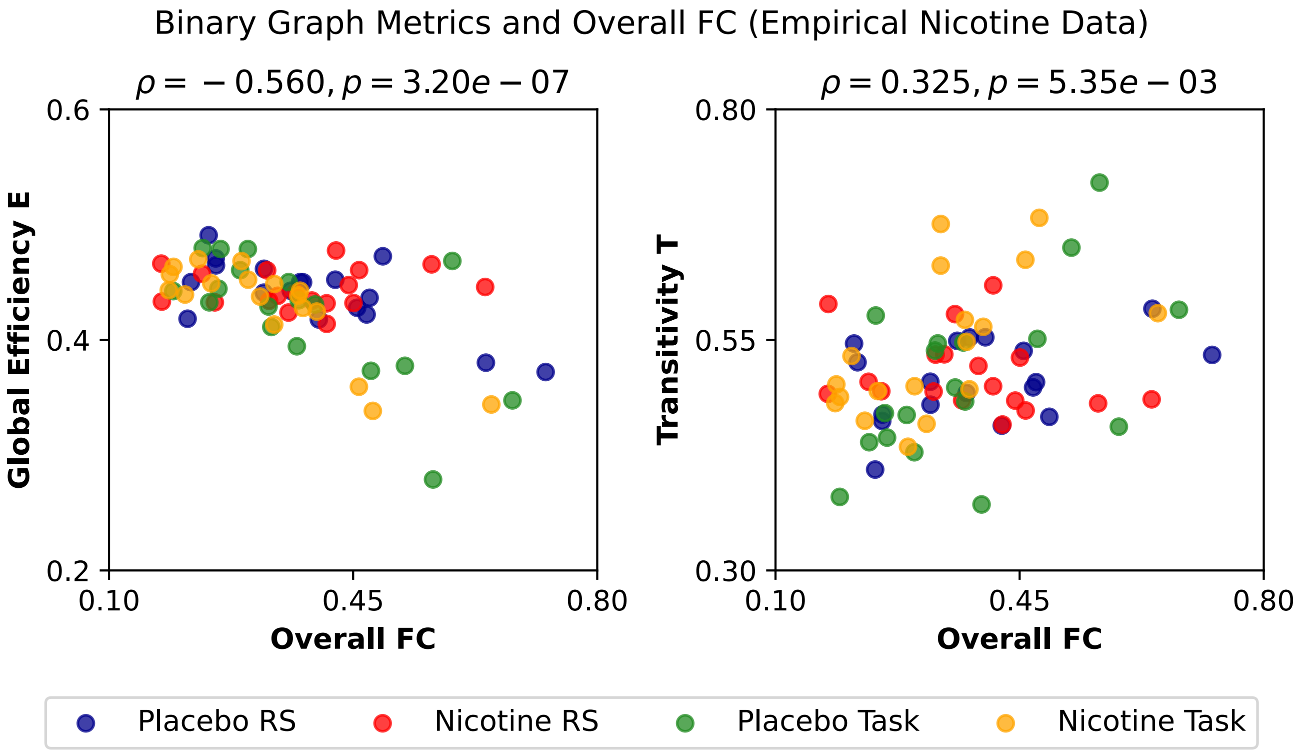


**Fig. S 2. Effect of overall FC on functional network metrics (empirical data).** Pearson's correlation $\rho$ was used as a correlation measure. Points correspond to participants (18 subjects per condition).

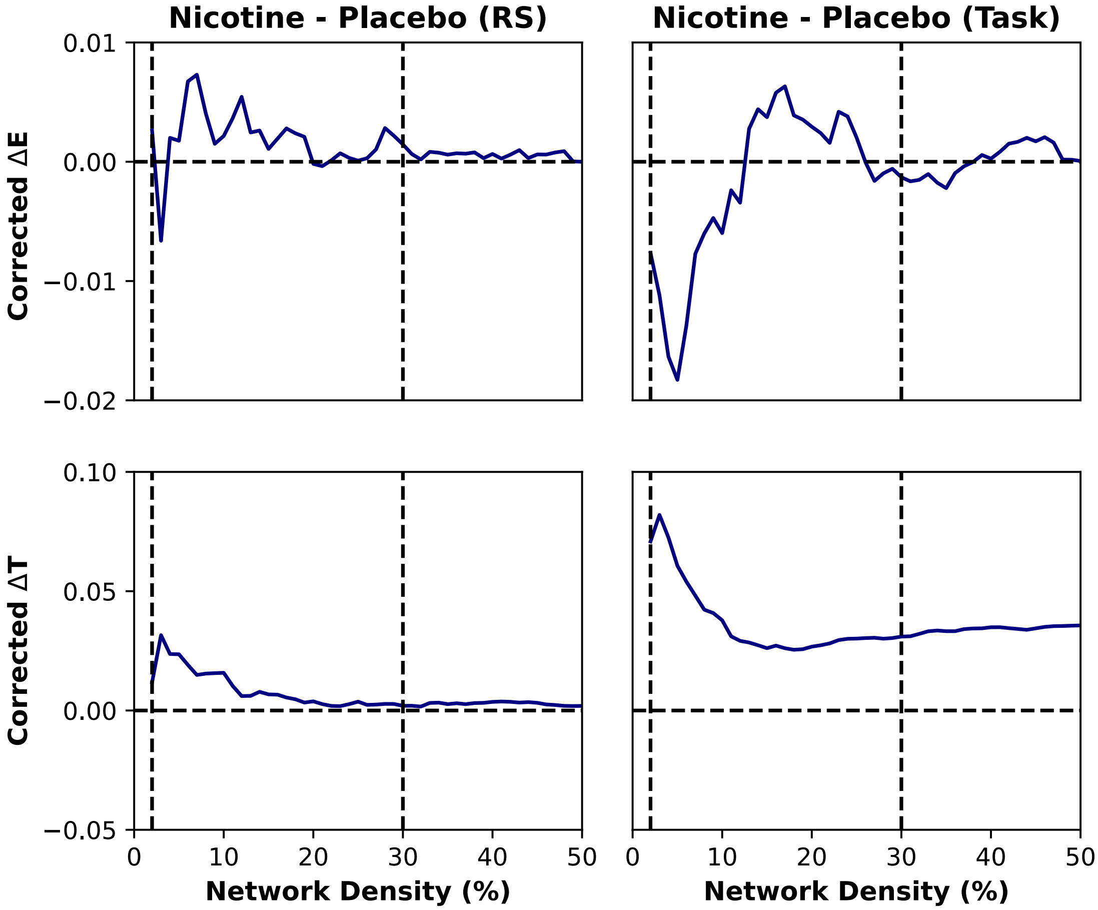


**Fig. S 3. Empirical graph metrics as functions of proportional thresholds.** Left and right columns correspond to resting-state (RS) and task blocks, respectively. Metrics (Transitivity, T, and Global Efficiency, E) were computed for a particular network density, and then corrected for overall FC. $\Delta$ denotes the difference nicotine minus placebo. Vertical dotted lines enclose the region used for computing the area under the curve (AUC) (2 and 30 %).

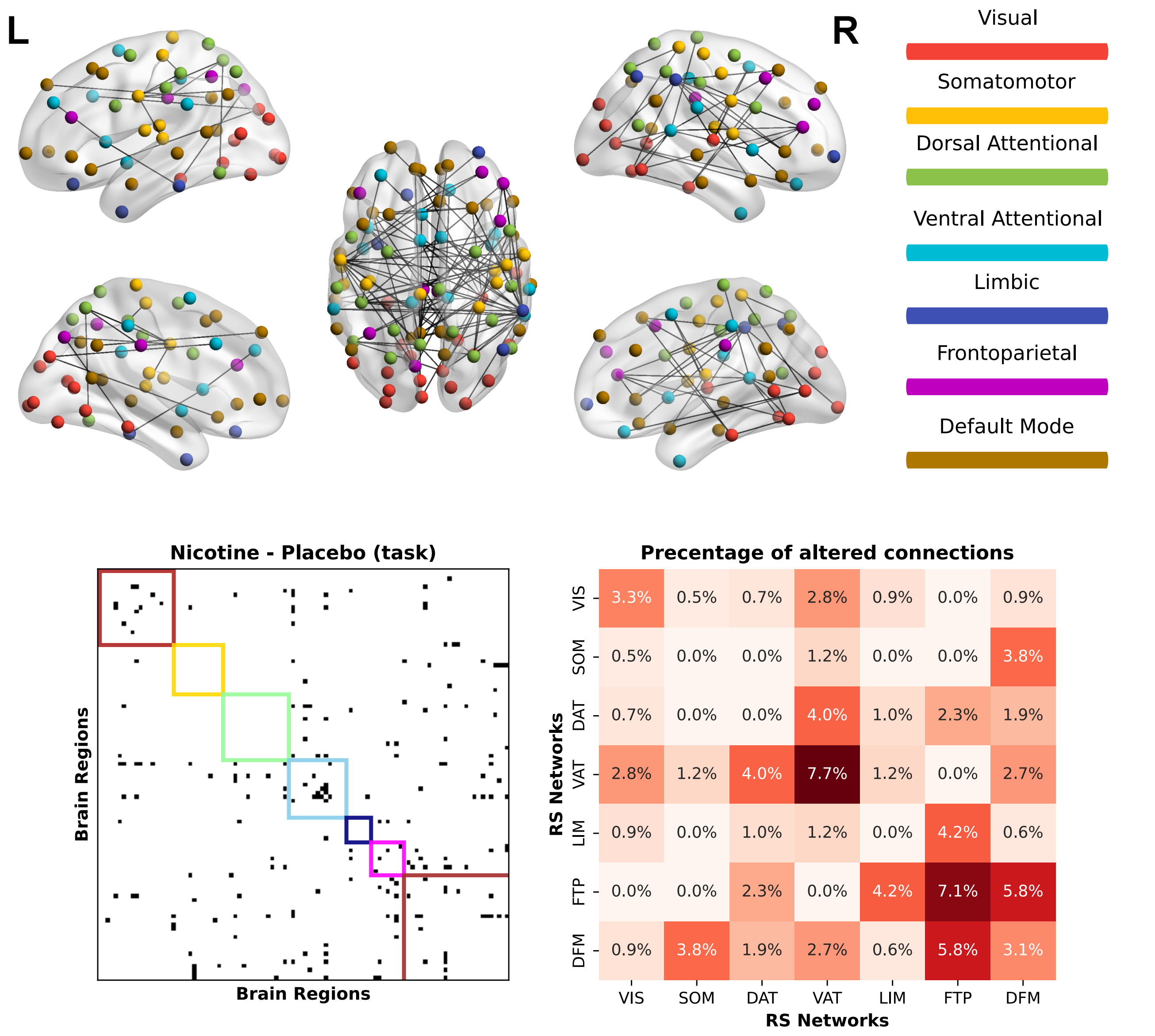


**Fig. S 4. Brain networks affected by nicotine during the task.** Colors represent 7 networks (subdivisions) of the Schaefer 100 cortical parcellation. Links correspond to the subnetwork, obtained through network based-statistic analysis, of links that decreased the most their strength by nicotine during the task. The subnetwork is also represented in the bottom-left matrix (black dots represent links). Regions within the matrix were ordered in accordance with their membership to the 7 networks considered in the parcellation; the colored squares represent each network. The percentage of altered connections, within and between networks, are depicted in the bottom-right matrix.

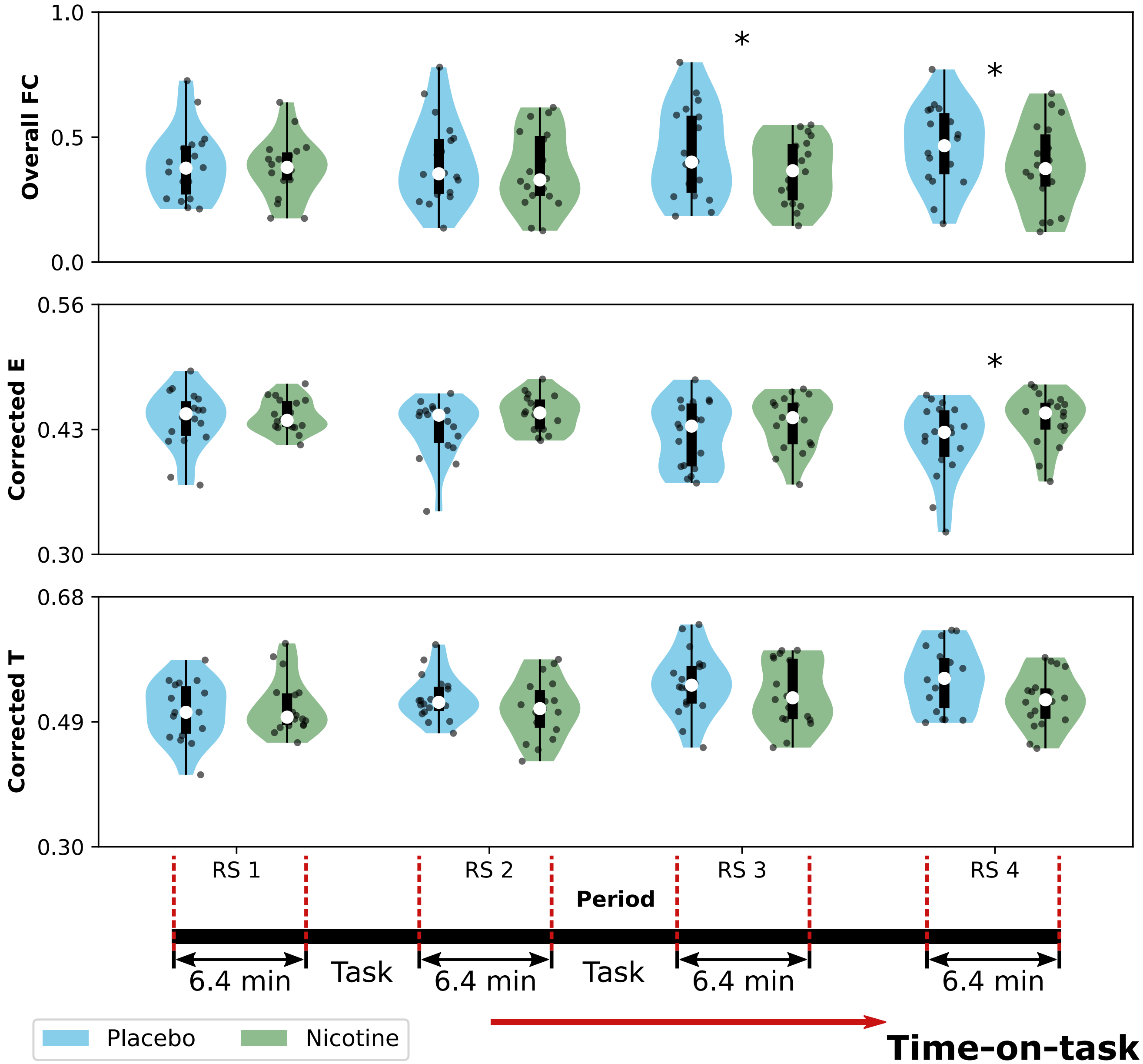


**Fig. S 5. Changes in functional connectivity across RS periods. A)** Global correlations (overall FC); **B)** corrected global efficiency $E$; and **C)** and corrected transitivity $T$. Violin plots represent the distributions of points (18 subjects per condition); the box plot is constituted by the median (white circle), the first and third quartiles (box limits), and sample range (whiskers). $*$ : $p < 0.05$.

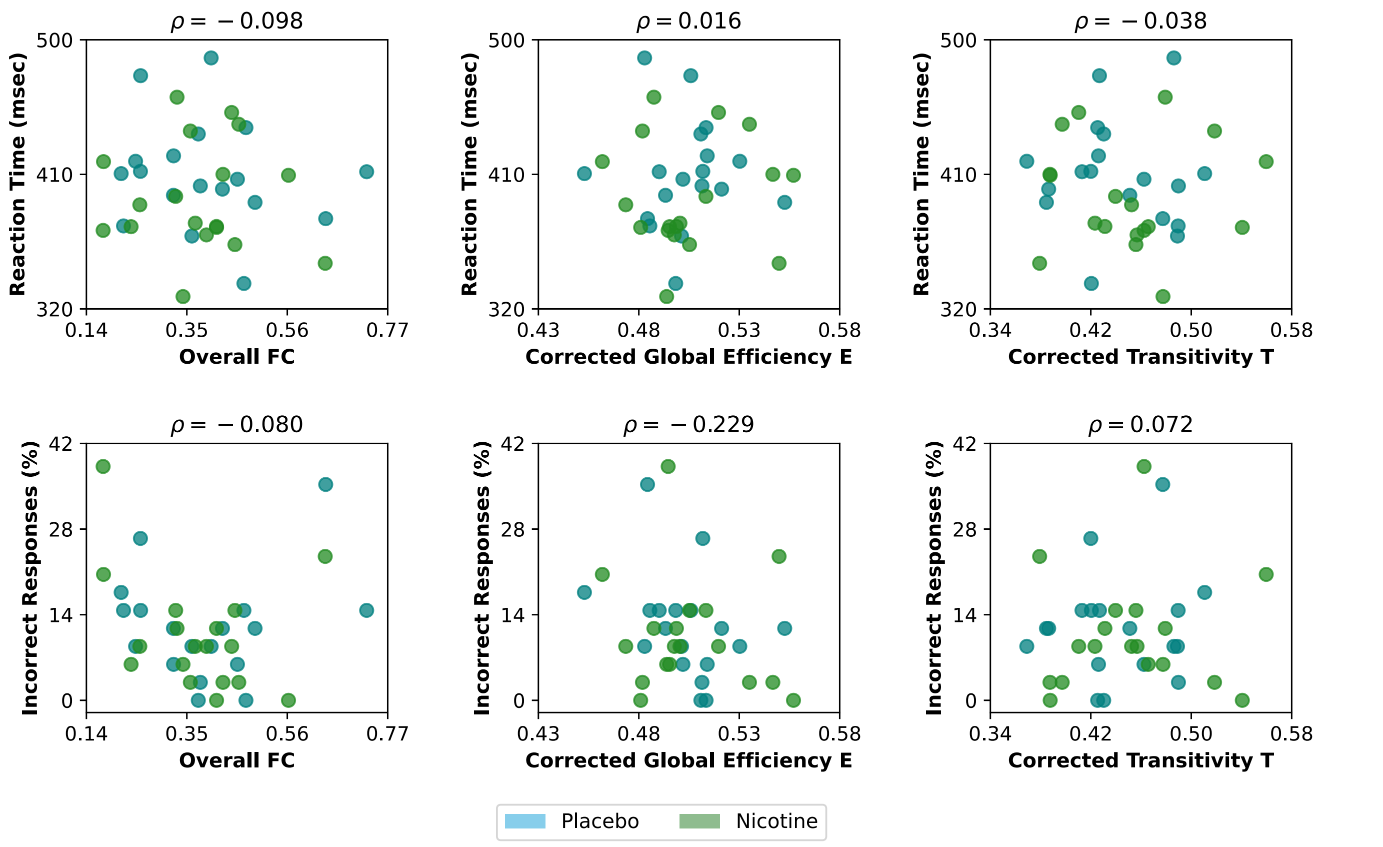


**Fig. S 6. Correlation of RS FC metrics with performance. A)** Correlation with the reaction time (RT) of the Go trials. **B)** Correlation with the percentage of incorrect responses of the No-Go trials. Pearson's $\rho$ was used as a correlation measure.

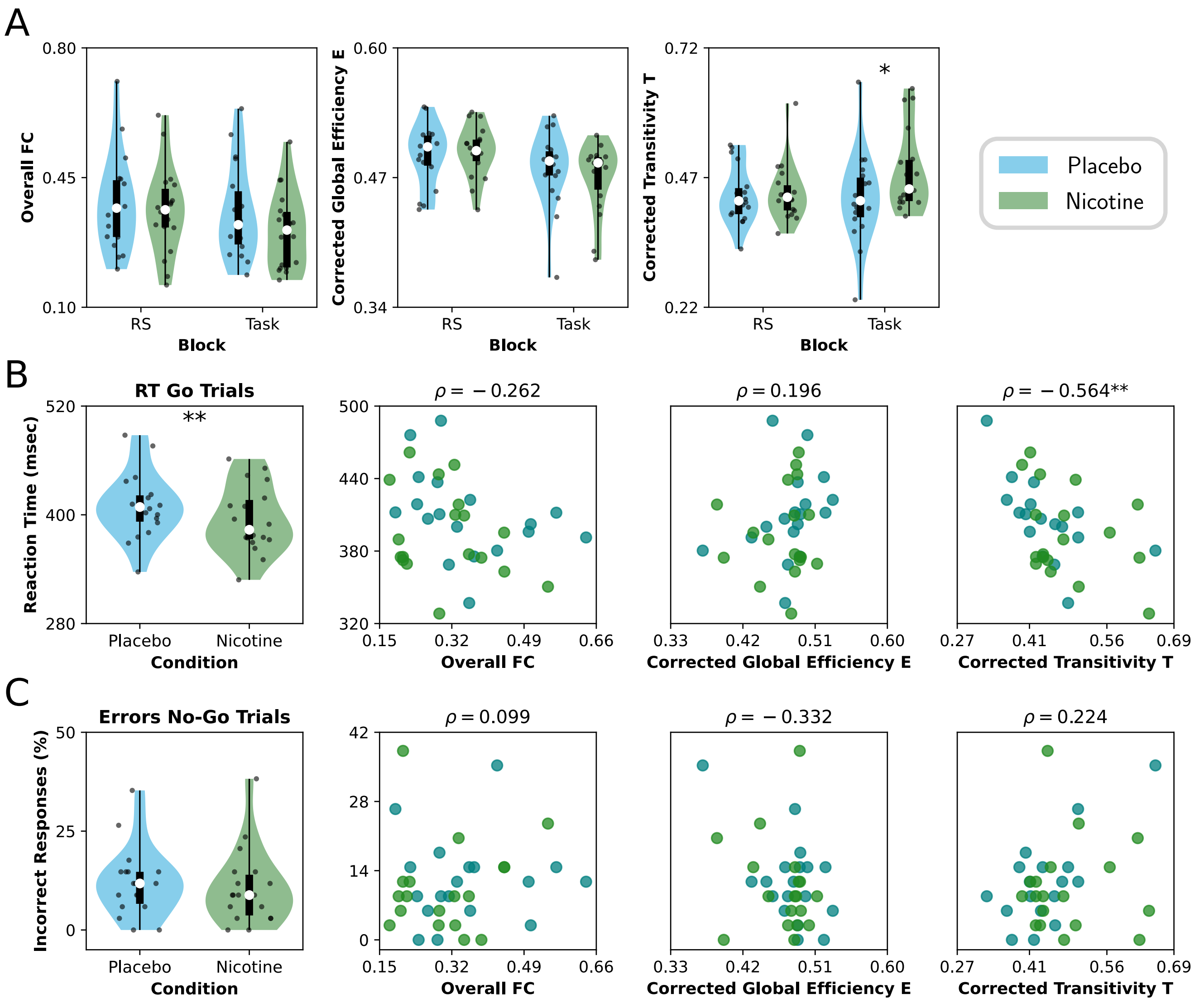


**Fig. S 7. Re-analysis of the empirical data using AAL90 parcellation. A)** Effect of nicotine on FC and functional network topology, both in RS and task. Violin plots represent the distributions of points (18 subjects per condition); the box plots represent the median (white circle), the first and third quartiles (box limits), and sample range (whiskers). **B)** Reaction time (RT) of the Go trials (left) and its correlation with FC metrics during the task. **C)** Percentage of incorrect responses of the No-Go trials (left) and correlation with FC metrics during the task. Pearson's $\rho$ was used as a correlation measure. Points correspond to participants (18 subjects per condition). $** : p < 0.01$, $* : p < 0.05$.

| Overall FC | | Global Efficiency | | Transitivity | |
|---|---|---|---|---|---|
| **Rest** | **Task** | **Rest** | **Task** | **Rest** | **Task** |
| $D = -0.150$ | $D = -0.426$ | $D = 0.115$ | $D = -0.241$ | $D = 0.102$ | $D = 0.631$ |
| $p = 0.249$ | $p = 0.122$ | $p = 0.349$ | $p = 0.349$ | $p = 0.325$ | $p = 0.045$ |

**Table S 1. Summary of placebo vs nicotine statistical tests (Supplementary Figure 7).** The table reports Cohen's D effect sizes and the corrected $p$ values (using FDR).

| Behavioral performance | | Correlations | | | | | |
|---|---|---|---|---|---|---|---|
| | | Overall FC | | Global Efficiency | | Transitivity | |
| **RT** | **Errors** | **RT** | **Errors** | **RT** | **Errors** | **RT** | **Errors** |
| $D = -0.41$ | $D = -0.14$ | $\rho = -0.262$ | $\rho = 0.099$ | $\rho = 0.196$ | $\rho = -0.332$ | $\rho = -0.564$ | $\rho = 0.224$ |
| $p = 0.004$ | $p = 0.321$ | $p = 0.367$ | $p = 0.680$ | $p = 0.506$ | $p = 0.264$ | $p = 0.002$ | $p = 0.283$ |

**Table S 2. Summary of correlations with performance (Supplementary Figure 7).** The table reports the Cohen's D effect sizes, Pearson's $\rho$ and the corrected $p$ values (using FDR).

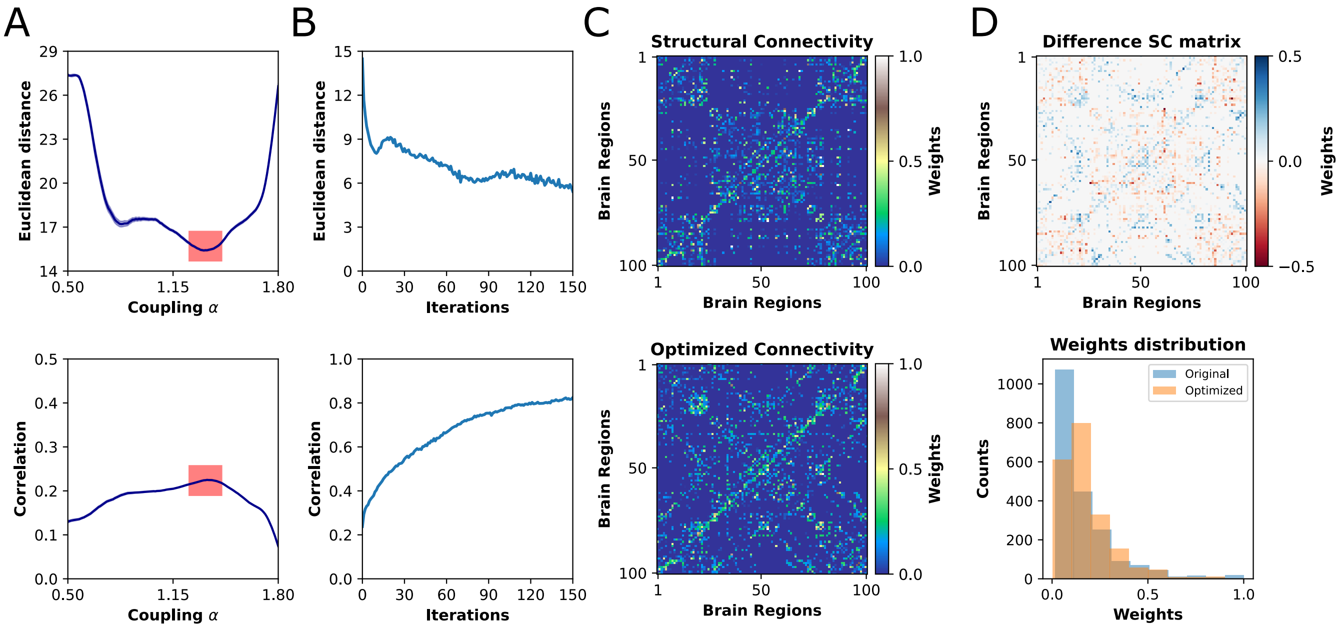


**Fig. S 8. Optimization of structural connectivity. A)** Fitting to placebo-RS condition using the original SC matrix. A better fit corresponds to low values of Euclidean distance and high values of Pearson's correlation. A value of $\alpha = 1.37$, within the red squares, was used for SC optimization. **B)** Euclidean distance and Pearson's correlation across iterations, for a fixed $\alpha = 1.37$. **C)** Original structural connectivity matrix (iteration number 0, top) and optimized matrix (final iteration, bottom). **D** Top, point-wise difference between the optimized and original SCs. Bottom, weights distributions of each SC.

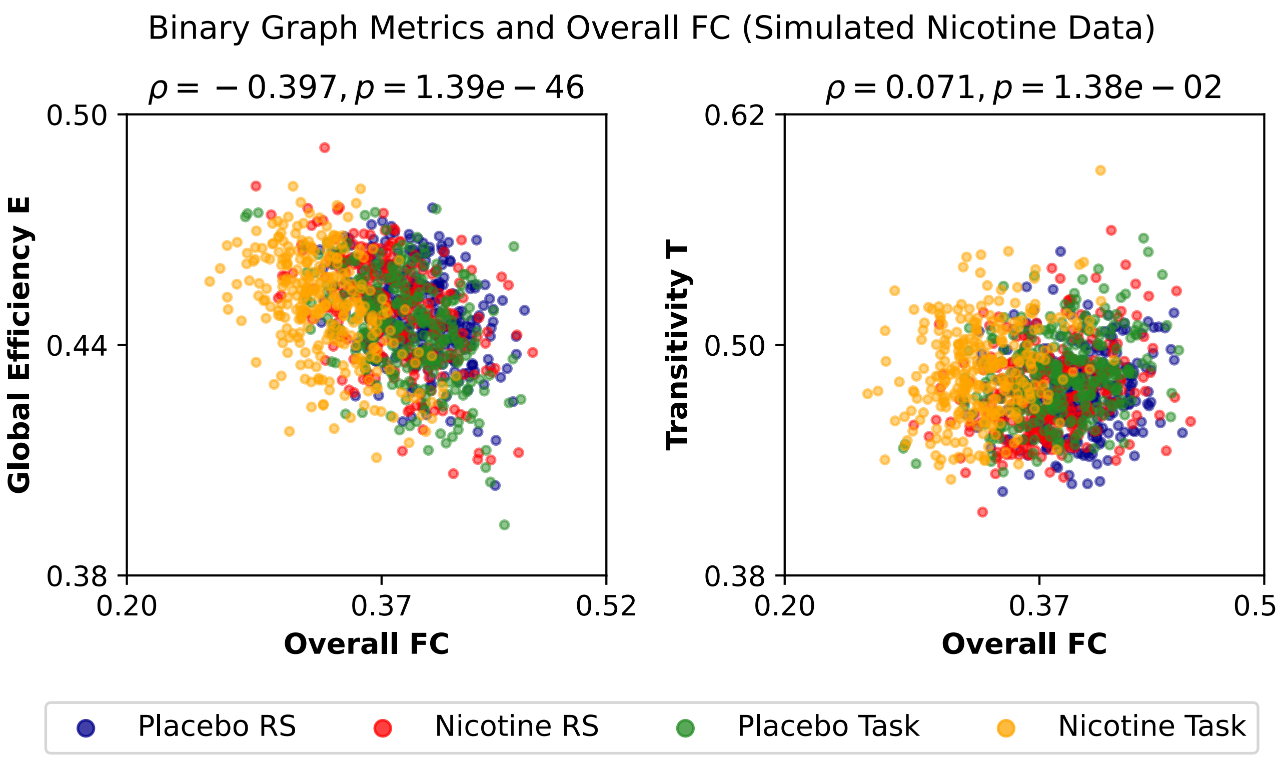


**Fig. S 9. Effect of overall FC on functional network metrics (simulated data).** Pearson's $\rho$ was used as a correlation measure. Points correspond to different random seeds (300 seeds per condition).

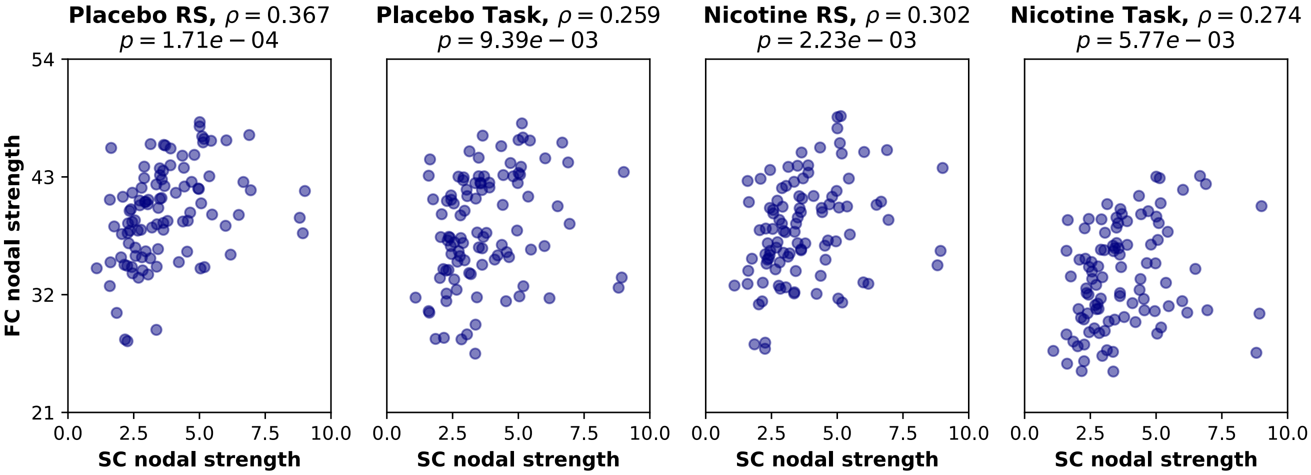


**Fig. S 10. Correlation between the empirical FC nodal strength (RS and task) with SC nodal strength.** Pearson's $\rho$ was used as a correlation measure. Points represent each of the Schaefer-100 brain regions. $** : p < 0.01$, $*** : p < 0.001$.
